# Engineering a SARS-CoV-2 vaccine targeting the RBD cryptic-face via immunofocusing

**DOI:** 10.1101/2024.06.05.597541

**Authors:** Theodora U.J. Bruun, Jonathan Do, Payton A.-B. Weidenbacher, Peter S. Kim

## Abstract

The receptor-binding domain (RBD) of the SARS-CoV-2 spike protein is the main target of neutralizing antibodies. Although they are infrequently elicited during infection or vaccination, antibodies that bind to the conformation-specific cryptic face of the RBD display remarkable breadth of binding and neutralization across *Sarbecoviruses*. Here, we employed the immunofocusing technique PMD (protect, modify, deprotect) to create RBD immunogens (PMD-RBD) specifically designed to focus the antibody response towards the cryptic-face epitope recognized by the broadly neutralizing antibody S2X259. Immunization with PMD-RBD antigens induced robust binding titers and broad neutralizing activity against homologous and heterologous *Sarbecovirus* strains. A serum-depletion assay provided direct evidence that PMD successfully skewed the polyclonal antibody response towards the cryptic face of the RBD. Our work demonstrates the ability of PMD to overcome immunodominance and refocus humoral immunity, with implications for the development of broader and more resilient vaccines against current and emerging viruses with pandemic potential.

**GRAPHICAL ABSTRACT:** 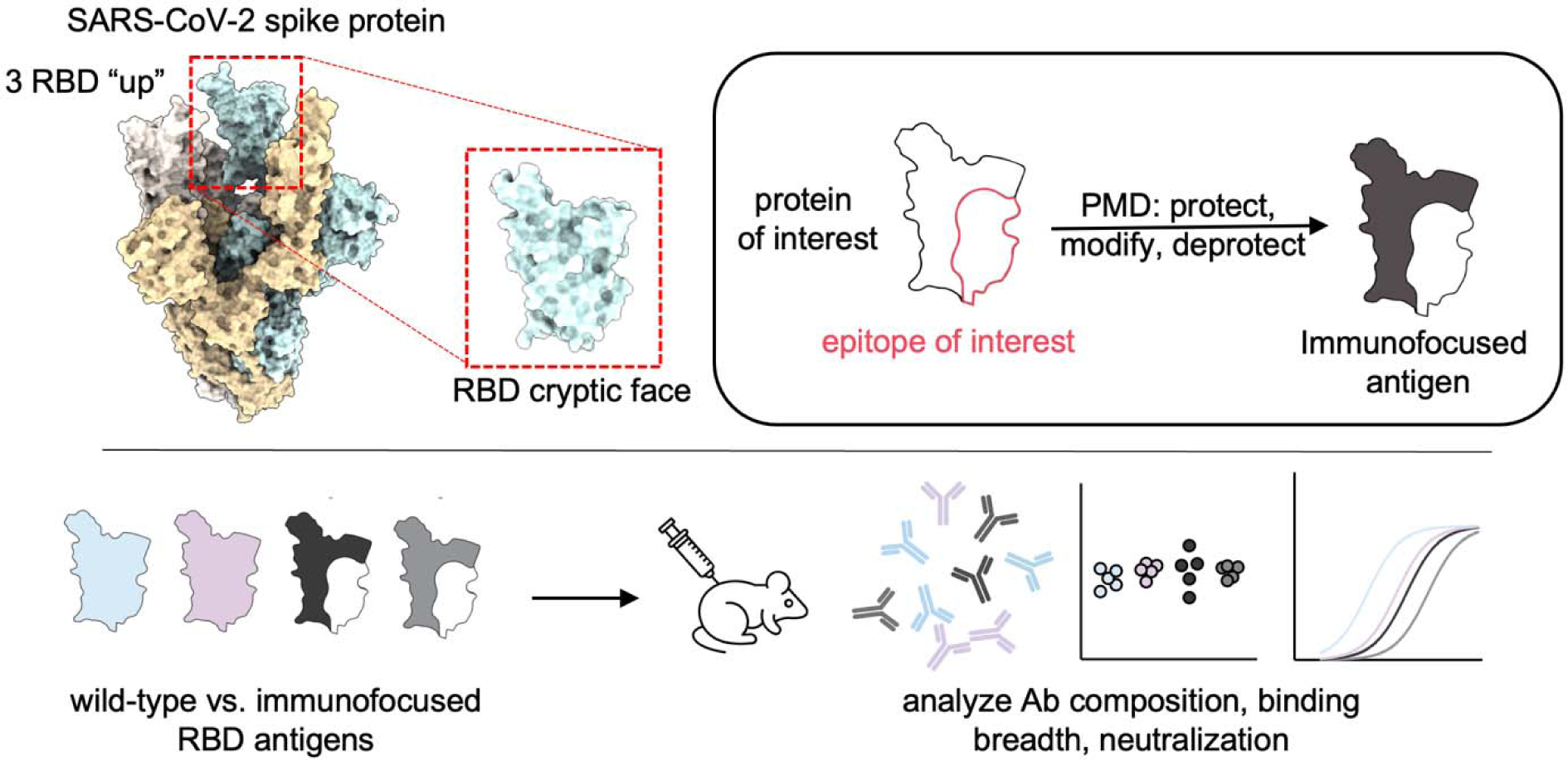

**SYNOPSIS:** Using the immunofocusing technique protect, modify, deprotect (PMD) we engineer SARS-CoV-2 receptor-binding domain (RBD) immunogens that generate broadly neutralizing antibodies against the conserved cryptic face. The ability to refocus humoral immunity to broadly conserved epitopes has important implications for the design of universal vaccines.

## INTRODUCTION

A major challenge in vaccinology is the ability to create vaccines that can provide protection against current as well as emerging variants of a given virus. The recent COVID-19 pandemic, caused by the SARS-CoV-2 *Sarbecovirus*, led to the development of numerous FDA-approved vaccines with unprecedent speed.^1–3^ However, the rapid evolution of SARS-CoV-2 variants of concern (VOCs) that are resistant to both infection and vaccine-elicited neutralizing antibodies has necessitated multiple monovalent and bivalent vaccine boosters.^4–6^ A vaccine that is robust to immune escape in the current SARS-CoV-2 pandemic as well as future potential *Sarbecovirus* spillover events is an important unmet public health need.^7,8^

One strategy to develop universally protective vaccines is to direct the humoral immune response towards on an antigenic region of a virus that is broadly conserved across viral variants and strains.^9^ This approach, termed immunofocusing, relies on protein engineering to diminish B-cell responses to non-neutralizing, immunodominant and subtype-specific epitopes while focusing the response towards the epitope of known broadly neutralizing antibodies. In the case of SARS-CoV-2, neutralizing antibodies have been isolated against four major areas of the spike protein: the N-terminal domain (NTD), receptor-binding domain (RBD), the S2 stem helix, and the S2 fusion peptide.^10^ However, the majority of potent and broadly neutralizing antibodies target the receptor-binding domain (RBD) which mediates attachment to human angiotensin converting-enzyme 2 (ACE2), the receptor for SARS-CoV-2 on human cells.^11–13^

Structural studies have found that RBD-binding antibodies can be broadly divided into four classes: those that bind the receptor-binding motif (RBM) and overlap with and block the ACE2-binding site (classes 1 and 2), those that bind the open face of the RBD which is accessible when the RBD is in the “up” or “down” conformation (class 3), and those that bind the cryptic face of the RBD, which is only accessible to antibodies in the “up” conformation (class 4).^14^

The class 4 cryptic epitope (also referred to as the RBD-6/RBD-7 site,^15^ the RBS-D/CR3022 site,^16,17^ the F2/F3 site,^18^ or the site II/core RBD^13^) is highly conserved across VOCs and distant *Sarbecovirus* strains.^17,19,20^ However, class 4 antibodies targeting the cryptic face make up a very small proportion of elicited antibodies post-vaccination or infection, suggesting that this site is subdominant to other portions of the RBD.^21–25^ Despite this, it has been possible to isolate rare monoclonal antibodies (mAbs) that bind to the cryptic face and show exquisite breadth of binding.^19,23,26–33^ Although cryptic-face mAbs are often less potent in neutralizing activity, there are examples of both broad and potent class 4 antibodies including S2X259,^34^ SA55,^18,35^ VacW-209,^36^ AGD20,^17^ DH1047^37^, and most recently SC27.^38^ These results suggest that if a vaccine could elicit antibodies that bind to the cryptic face of the RBD, it might be possible to create a pan-*Sarbecovirus* vaccine.

Previous efforts to create an immunofocused-RBD vaccine have utilized techniques such as epitope dissection,^39,40^ mosaic display,^41,42^ epitope masking via glycosylation,^43–47^ and cross-strain boosting.^48–50^ In this work, we employ the immunofocusing technique called protect, modify, deprotect (PMD)^51^ to create a cryptic face-targeting RBD vaccine. In PMD, a target epitope is protected by binding to a mAb, after which lysines outside of the antibody-antigen interface are covalently modified through reaction with *N*-hydroxysuccinimide (NHS) ester-modified polyethylene glycol (PEG) moieties. Following dissociation of the mAb, the installed PEG chains decrease the immunogenicity of the resulting immunogen at epitopes outside of the desired epitope.

To generate a cryptic face-targeting RBD vaccine using PMD, we used the broadly binding and neutralizing class 4 mAb S2X259 as our protecting mAb.^34^ To install PEG chains on the remainder of the RBD surface not shielded by S2X259, we engineered an RBD variant that contained eleven additional lysines in the epitopes of off-target class 1,2,3 antibodies. In addition, we used PEG conjugated to either NHS-esters or Bis-NHS-esters to generate our immunofocused PMD-RBD antigens.

While both wild-type RBD and PMD-RBD antigens generated high antibody titers in immunized mice against a panel of *Sarbecoviruses*, compared to the unmodified RBD the PMD-RBD antigens induced a more consistent neutralizing activity against the more divergent *Sarbecoviruses* SARS-CoV-1 and WIV-1.

To directly assess the ability of PMD to focus the B-cell response specifically towards the RBD’s cryptic face, we developed and validated an assay that uses an RBD epitope knock-out protein to measure the proportion of class 4 antibodies present in mouse polyclonal antisera after vaccination with wild-type RBD and two PMD-RBD antigens. Using this assay, we selectively depleted mouse antisera of RBD-directed antibodies except those binding the cryptic face and then tested the ability of the depleted antisera to bind and neutralize SARS-CoV-2. We found that PMD-RBD antigens elicited a higher proportion of class 4 antibodies compared to wild-type RBD and that these antibodies are capable of neutralizing SARS-CoV-2 and SARS-CoV-2 Beta. To our knowledge, this is the first direct evidence that immunofocusing is able to alter the composition of polyclonal antibodies elicited in a vaccine context. Our results demonstrate the generalizability of PMD as an immunofocusing tool to create broad-spectrum vaccines. Furthermore, we expand upon previous serum depletion assays^11,52,53^ by using an RBD epitope knock-out protein to assess the composition of antibodies in a polyclonal mixture. This assay will be useful in the assessment and comparison of future immunofocused vaccines.

## RESULTS

### Engineering the SARS-CoV-2 RBD for PMD

To express the RBD of SARS-CoV-2 in mammalian (Expi-293F) cells, we initially codon-optimized and cloned the sequence corresponding to residues 319-541 from the SARS-CoV-2 Wuhan Hu-1 receptor-binding domain (RBD) into the pADD2 vector. Initial purification and SDS-PAGE characterization of the product revealed a protein with a tendency to dimerize (**Fig. S1**). By removing eight residues at the C-terminal end of the RBD sequence that included one cysteine (leaving residues 534-541), we could produce a monomeric RBD protein in high yield (**Fig. S1**).

To validate the proper folding of our RBD protein, we used bio-layer interferometry (BLI) to perform a binding competition assay with four antibodies representing the major epitopes on the surface of RBD (class 1: CB6,^54^ class 2: C002,^14^ class 3: S309,^55^ class 4: S2X259) (**Fig.1a**). As expected, RBD-binding antibodies CB6 and C002 competed with each other when bound to RBD, while CB6 also competed with class 4 S2X259 due to partial overlap with the cryptic face epitope (**Fig.1b**). S309, which binds the open face of the RBD, did not compete with any of the other antibodies.

Since we were interested in immunofocusing to the highly conserved class 4 cryptic face epitope we deployed S2X259 as the protecting mAb for PMD. Unlike many other class 4 mAbs, S2X259 uses both its VH and VL domains to make extensive contacts with the RBD and has a large binding footprint (**Fig. 1a**). Based on the structure of RBD (PDB ID: 7M7W), we identified regions lacking surface-exposed lysines (i.e., “antigenic holes”) that would retain their antigenicity after the PMD protocol. By analyzing data from a deep mutational-scanning library of the RBD,^57^ we initially selected 11 amino acids within the antigenic holes to screen for lysine installation in addition to the 10 naturally occurring lysines on the surface of RBD (**Fig. 1c**). We therefore created 11 individual lysine variants via site-directed mutagenesis and analyzed their expression compared to wild-type RBD (RWT) using dot-blots (**Fig. S2**). Seven of these variants retained >40% expression compared to wild-type RBD, three retained >30% expression, and one variant completely ablated expression (**Fig. 1d**). Based on lysine location, we cloned R346K, V367K, N440K, R466K, T478K, F490K, and L518K into RWT to create an RBD containing seven additional lysines (R7K). Compared to RWT, R7K was less thermostable (melting temperature of 46°C, compared to 53°C for RWT) (**Fig. 1e**). Since R7K still retained good expression (>20 mg/L), we performed another round of screening to increase the density of lysines within the epitopes of class 1,2,3 antibodies. We tested an additional 10 surface-exposed lysine residues in the context of R7K (**Fig. 1f**). We took the four or five best-expressing variants and cloned them into the R7K plasmid to create R11K and R12K, respectively. Since we found that cloning additional lysines lowered the melting temperature of R7K, we also added known RBD repackaging substitutions Y365F, F392W, and V395I to improve stability.^56^ We successfully expressed and purified R11K and R12K and found that both had similar thermal melting profiles and melting temperatures as RWT (**Fig.1g**). The expression level of the R11K protein was higher than that of R12K, so we chose R11K as the final candidate for PMD. To ensure that the introduction of 11 lysines did not affect the S2X259 epitope, we used BLI to compare binding of the fragment antigen-binding region (Fab) of S2X259 to RWT and R11K (**Fig.1h**). The binding profile of S2X259 Fab and the calculated *K*_D_s (0.97 ± 0.5 nM for RWT and 1.5 ± 0.3 nM for R11K) confirmed the integrity of the S2X259 conformational epitope in R11K.

**Figure 1.**
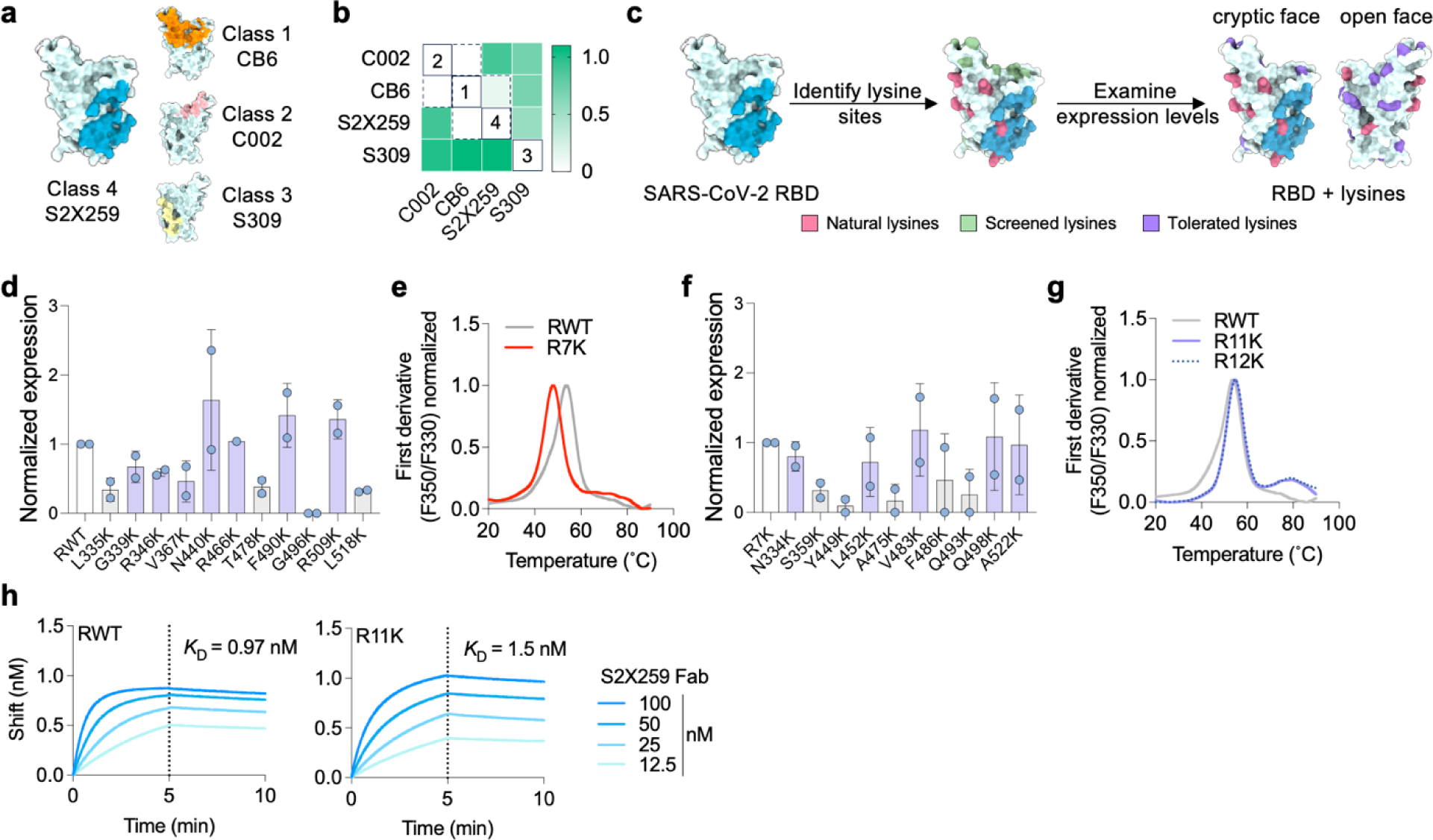
Engineering the RBD of SARS-CoV-2 for PMD. **a,** Structure of SARS-CoV-2 RBD (PDB ID: 7M7W) showing the binding footprint of one representative antibody from each of the four classes of RBD-binding antibodies; class 1 (Ab CB6, orange), class 2 (Ab C002, light pink), class 3 (S309, yellow), and class 4 (S2X259, blue). **b**, BLI competition binding assay of RBD-directed antibodies binding to RBD. Loaded antibodies are displayed in rows and competing antibodies are displayed in columns. White color indicates no binding of the tested anitbody, indicating that the antibodies compete for binding (all competing antibodies are enclosed within dotted lines, unique competition groups are enclosed within solid lines). **c**, Screening and identification of permissive lysine installations on RBD. In addition to 10 natural lysines (pink), 11 single lysine substitutions (green) were introduced by site-directed mutagenesis and all 12 proteins were transiently expressed. **d**, Expression of lysine-substituted RBDs in Expi-293F cells quantified by dot-blot and normalized to the expression level of wild-type RBD (RWT). Data are presented as mean ± standard deviation (*n* = 2 biological replicates). Lysines with >40% expression compared to RWT are colored purple in **c**. **e**, Thermal melting profile of RWT and R7K (RWT with additional R346K, V367K, N440K, R466K, T478K, F490K, L518K substitutions) measured by differential scanning fluorimetry. **f**, Expression level of single lysine substitions cloned into the R7K protein. Expression level normalized to that of R7K. Data are presented as mean ± standard deviation (*n* = 2). **g**, Thermal melting profile of RWT, R11K (R7K plus L452K, V483K, Q498K, and A522K), and R12K (R7K plus N334K, L452K, V483K, Q498K, and A522K), measured by differential scanning fluorimetry. R11K and R12K also contain the repackaging substitutions Y365F, F392W, V395I to improve stability.^56^ **h**, BLI binding curves of RWT and R11K with the fragment antigen-binding (Fab) of S2X259. Association was monitored for 5 min (dotted lines), after which dissociation was monitored for 5 min.

### Expression, characterization, and antigenicity of immunofocused RBD antigens using PMD

To produce immunofocused RBD antigens using PMD, we started by reacting purified S2X259 IgG with AminoLink^TM^ agarose beads to create S2X259-conjugated resin. After protection of R11K by binding to the S2X259-conjugated resin, we modified the RBD by reaction of surface-exposed lysines with NHS-esters conjugated to PEG chains (**Fig. 2a**). The initial protection step was necessary because the S2X259 epitope contains a lysine (K378) which is reactive to NHS-esters and modification with NHS-esters containing PEG blocks binding of S2X259 (**Fig. S3**). K378 is also highly conserved (found in 46 of 52 SARS CoV homologs)^58^ and could be important for the elicitation of class 4 antibodies.

**Figure 2.**
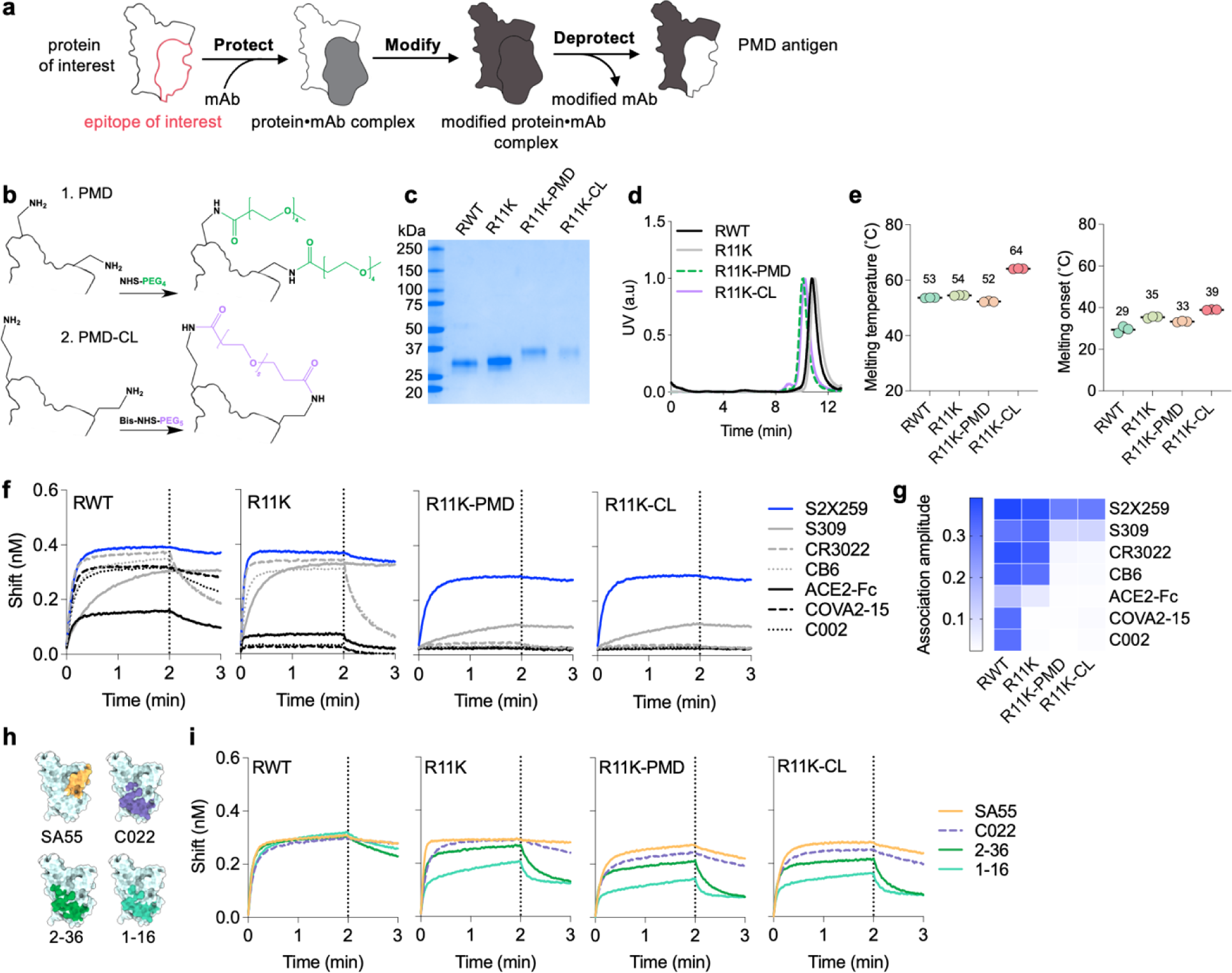
Using PMD to create immunofocused antigens. **a**, Schematic depicting the PMD strategy. First, RBD is bound to S2X259-conjugated resin (protect). Then, the surface of the protein-mAb complex is rendered non-immunogenic through chemical conjugation (modify). In the last step, the S2X259 epitope is exposed (deprotected) by dissociating it from the mAb resin. **b**, Illustration of the modification of two surface exposed lysines on R11K with NHS-PEG_4_ as in PMD (top), or the cross-linking of two surface exposed lysines with BIS-NHS-PEG_5_ as in PMD-CL (bottom). **c**, SDS-PAGE analysis of the four antigens after expression and purification. **d**, Normalized size-exclusion chromatographic traces of purified proteins. **e**, Thermal melting temperature and onset temperature of melting for RWT, R11K, R11K-PMD, and R11K-CL measured by differential scanning fluorimetry. Data are presented as mean ± standard deviation (*n* = 3 replicates). **f**, BLI binding curves of RBD-directed antibodies from all four classes and ACE2 (expressed as a dimer due to fusion to a human Fc domain) to RWT, R11K, R11K-PMD, and R11K-CL. Association was monitored for 2 min (dotted lines), after which dissociation was monitored for 1 min. **g**, BLI association amplitude of each antibody binding to the four antigens (corresponding to curves shown in **f**). **h**, Epitope footprints of S2X259-like class 4 antibodies shown in color on the light blue SARS-CoV-2 RBD structure (PDB ID: 7M7W). **i**, BLI binding curves of S2X259-like antibodies shown in **h** binding to RWT, R11K, R11K-PMD and R11K-CL.

We created two PMD antigens to bias immune responses towards the RBD cryptic-face epitope, R11K-PMD and R11K-CL, by using either NHS-PEG_4_-methyl (NHS-PEG_4_, with four ethylene glycol units) or Bis-NHS-PEG_5_ (a PEG_5_ linker containing an NHS ester on each end), respectively, in the modification step (**Fig. 2b**). The use of a Bis-NHS-ester allows for neighboring lysines to become cross-linked (CL), with the goal of imparting additional stability and minimal disruption to the conformation of the underlying protein.

Following the deprotection step of PMD, in which R11K-PMD or R11K-CL were dissociated from the S2X259-resin using low pH, the immunofocused antigens were purified to homogeneity and analyzed via gel electrophoresis (**Fig. 2c,d**). As expected, R11K-PMD and R11K-CL had a higher molecular weight compared to RWT and R11K, reflecting the addition of the PEG moieties (**Fig. 2c,d**). As hypothesized, the cross-linking of lysines on the surface of R11K-CL imparted a stability benefit which increased the melting temperature of R11K-CL by 10°C compared to R11K and melting onset of 4°C as compared to R11K (**Fig. 2e**). In contrast, the conjugation of PEG_4_ to R11K to generate R11K-PMD resulted in a 2°C decrease in both melting temperature and melting onset compared to R11K (**Fig. 2e**).

Using BLI, we compared the binding of a panel of six antibodies and ACE2 to RWT, R11K, R11K-PMD and R11K-CL (**Fig. 2f,g**). The antibody panel represented all classes of RBD-directed antibodies; class 1 (CB6^54^), class 2 (C002^14^ and COVA2-15^59^), class 3 (S309^55^), and class 4 (S2X259 and CR3022^29^) and was thus able to assess the antigenicity of each RBD antigen. As expected, all six antibodies and ACE2 (expressed as a dimer due to fusion to a human Fc domain) were able to bind RWT, while the addition of lysines in R11K disrupted the binding of class 2 antibodies primarily. In contrast, addition of PEG to the surface of R11K-PMD and R11K-CL completely diminished binding of all antibodies except S2X259, and to some extent antibody S309 which showed minor binding (**Fig.2 f,g**). Although increasing the length of PEG conjugated to R11K to greater than the original *n*=4 construct could completely ablate binding of S309 (**Fig. S4a-e**), a small-scale immunization trial in mice with R11K antigens conjugated to NHS-PEG_n_ reagents (*n* = 2,4,8,12, or 24) showed that increasing PEG length correlates with reduced immunogenicity (**Fig. S4f-h**). Thus, we chose to proceed with R11K conjugated with NHS-PEG_4_ (R11K-PMD) and Bis-NHS-PEG_5_ (R11K-CL) despite their both showing minor binding to the class 3 antibody.

To confirm that the addition of PEG to R11K-PMD and R11K-CL did not affect binding by other class 4 antibodies, we assessed the binding of SA55 (defined by PDB ID: 7Y0W),^18^ C022 (PDB ID: 7RKU),^19^ 2-36 (PDB ID: 7N5H),^26^ and 1-16 (PDB ID: 7JMW)^28^ (**Fig. 2h**) to all four antigens – RWT, R11K, and the two PMD antigens. Binding of all S2X259-like class 4 antibodies remained intact in R11K-PMD and R11K-CL (**Fig. 2i**), confirming that the additional 11 surface-exposed lysines on the RBD surface did not disrupt the structure of the S2X259 epitope, and the addition of PEG via PMD/CL decreased antigenicity at off-target sites while maintaining on-target antigenicity.

### Immunogenicity of wild-type and immunofocused RBD antigens

To investigate the immunogenicity of the RBD antigens, we immunized four groups of mice with RWT, R11K, R11K-PMD, or R11K-CL adjuvanted with CpG and alum (**Fig. 3a**). Mice were boosted three times with the same formulation at 5-week intervals to promote a strong immune response. We analyzed IgG titers to RBDs from five different clades within the *Sarbecovirus* subgenus one-week post-boost 2 (Week 11) and found that antibody titers to all five RBDs were high with no significant differences between the immunization groups (**Fig. 3b**). Although the titers against wild-type SARS-CoV-2 RBD were slightly lower for R11K-PMD and R11K-CL compared to RWT and R11K, the average IgG titer against RBDs from more distant clades (BtKY72, WIV-1, SARS-CoV-1, BM-4831) were either the same or higher for mice immunized with R11K-PMD and R11K-CL compared to RWT and R11K. In addition, an antibody-avidity ELISA assay^60^ indicated that more higher-avidity antibodies were generated by immunization with R11K-PMD and R11K-CL, with the most higher-avidity antibodies generated in the R11K-PMD-immunized mice (**Fig. S5a,b**).

**Figure 3.**
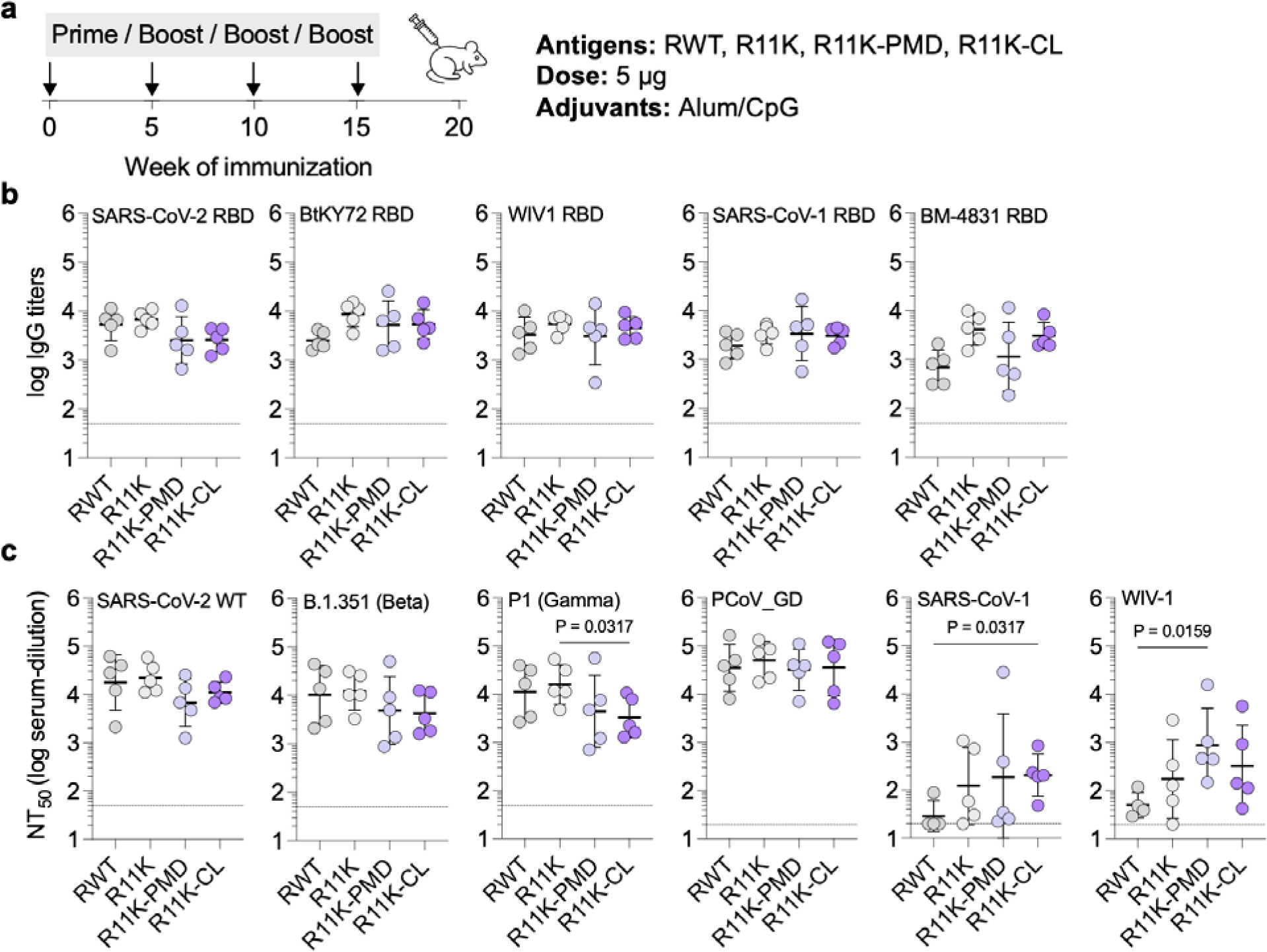
Immunogenicity of wild-type RBD and immunofocused RBD antigens. **a**, Schematic of the mouse immunization with a four-dose regimen occuring on days 0, 35, 70, and 105. Mice were immunized with 5 *ug* of antigen adjuvanted with alum/CpG (500 *u*g/20 *u*g) *via* intramuscular injection (*n* = 5 per group). **b**, Serum IgG titers on day 77 against biotinylated-RBDs from SARS-CoV-2 as well as more distantly related *Sarbecoviruses* including bio-BtKY72, bio-WIV-1, bio-SARS-CoV-1, and bio-BM-4831. **c**, Neutralization titers (NT_50_ – the serum dilution required to neutralize 50% of a given pseudotyped lentivirus) of day 112 serum against wild-type SARS-CoV-2 (D614G), the Beta and Gamma variants of SARS-CoV-2, and more distant SARS-CoV-1, WIV-1, and pangolin coronavirus PCoV_GD. Data are presented as geometric mean ± s.d. of the loglJtransformed values. Each circle represents a single mouse. Horizontal dotted lines indicate the limit of quantitation. Comparisons of two groups were performed using the two-tailed Mann-Whitney U test. *P* values of 0.05 or less were considered significant and indicated.

To assess broad neutralizing activity, we generated six *Sarbecovirus* spike-pseudotyped lentiviruses and performed neutralization assays with our mouse antisera in HeLa cells over-expressing ACE2 and the SARS-CoV-2 protease TMPRSS2. Neutralization titers were high for all immunization groups against SARS-CoV-2 (D614G) and the SARS-CoV-2 Beta VOC (**Fig. 3c, Fig. S6a,b**). Neutralizing titers against the SARS-CoV-2 Gamma VOC were high in all groups but significantly lower in the R11K-CL immunized mice (**Fig. 3c, Fig. S6c**). However, the neutralizing titers against lentiviruses from more-divergent *Sarbecovirus* clades were either the same across the immunization groups (as seen with the PCoV_GD lentivirus, **Fig. 3c, Fig. S6d**), or significantly higher in the R11K-PMD and R11K-CL immunized groups (as seen with the SARS-CoV-1 and WIV-1 lentiviruses) (**Fig. 3c, Fig. S6e,f**). In particular, the neutralization response in the R11K-CL immunized mice was most consistent, with all 5 mice in the immunization groups developing neutralization titers against SARS-CoV-1 and WIV-1.

### Assessing the quantity of class 4 antibodies in mouse polyclonal antisera

To quantify the proportion of S2X259-like antibodies elicited by our immunofocused antigens we first performed a competition ELISA in which biotinylated RBD bound to streptavidin coated ELISA plates were pre-incubated with either S309 antibody (class 3) or S2X259 antibody (class 4) before the addition of mouse antisera (**Fig. 4a**). In all immunization groups, competition with S309 led to only a small decrease in binding titers, between a one-to two-fold decrease. Similarly, competition with S2X259 showed comparable decreases in binding titers in the mice immunized with RWT (2.3-fold) and R11K (2.1-fold). In contrast, competition with S2X259 led to a 6-fold and almost 5-fold decrease in average binding titer of the R11K-PMD and R11K-CL immunized mice, respectively. This suggests that, consistent with their design, a much higher percentage of the measured binding titer in the groups immunized with the immunofocused antigens were due to antibodies targeting the class 4 S2X259-binding epitope.

**Figure 4.**
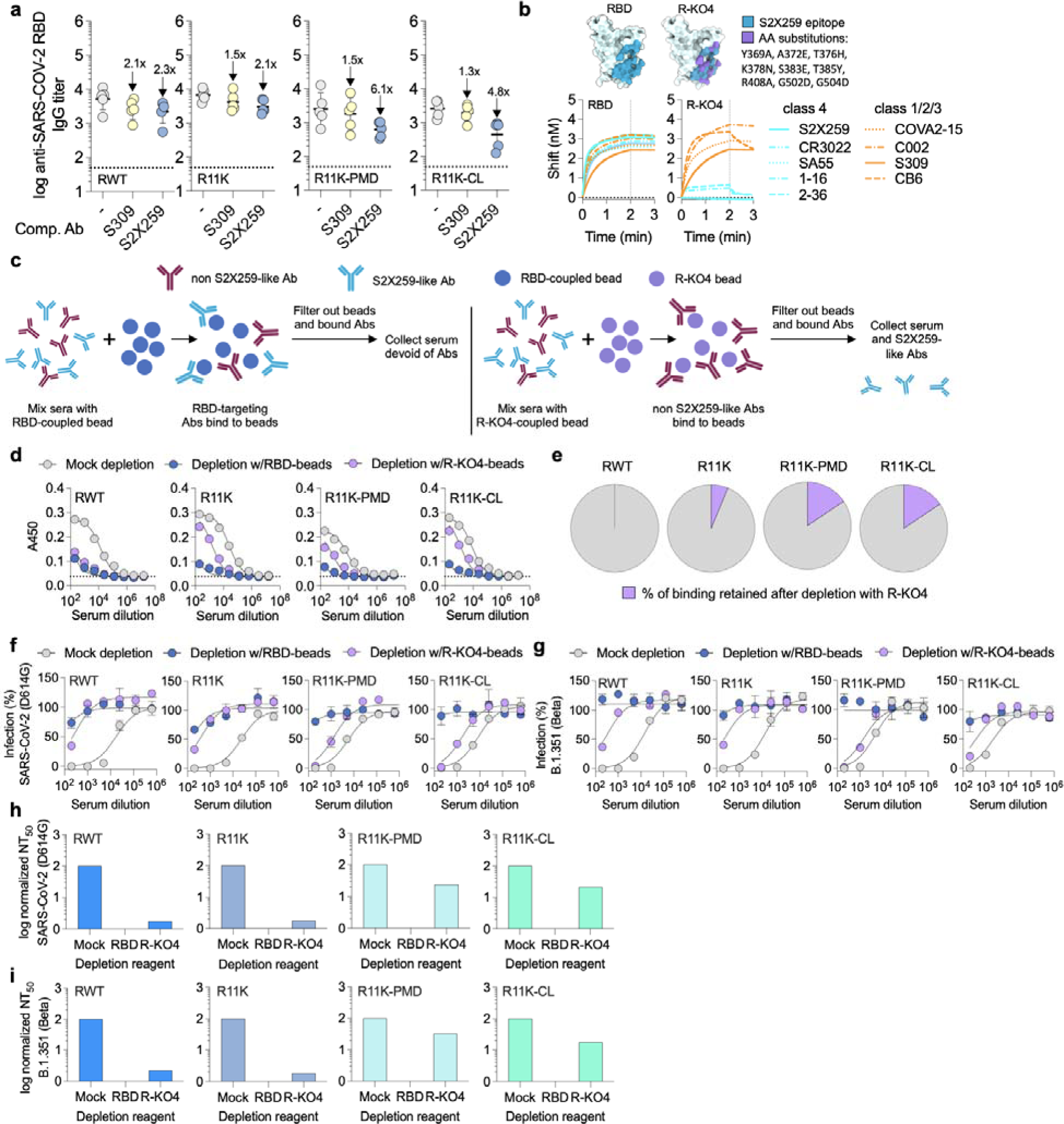
Quantifying the proportion of cryptic-face targeting antibodies elicited by immunization. **a**, Serum IgG binding titers of day-77 antisera to SARS-CoV-2 RBD in the presence of competing mAbs. Data are presented as geometric mean ± s.d. of the loglJtransformed values. Fold-change is indicated by arrows with numbers. Horizontal dotted lines indicate the limit of quantitation. **b**, Substitutions in the S2X259 epitope of SARS-CoV-2 RBD (PDB ID: 7M7W) to create R-KO4. Binding of class 4 and non-class 4 antibodies to RBD and R-KO4 measured by BLI. Association was monitored for 2 min (dotted lines), after which dissociation was monitored for 1 min. **c**, Schematic depicting the serum-depletion assay used to quantify the S2X259-like antibodies elicited by immunization. First, pooled antisera from immunized mice are mixed with neutravidin beads bound to either biotinylated RBD or biotinylated R-KO4. After incubation for 30 min, antibody-bound beads are filtered out, leaving antisera containing no RBD-directed antibodies, or only S2X259-like antibodies unable to bind to R-KO4. **d**, ELISA was used to measure binding of pooled antisera to SARS-CoV-2 RBD after depletion with either unbound beads (mock depleted serum), beads coupled to RBD (RBD-depleted serum), or beads coupled to R-KO4 (R-KO4-depleted serum). **e**, Percentage of binding retained after depletion with R-KO4, calculated by taking the EC_50_ (serum dilution at which half-maximal binding is achieved) of R-KO4-depleted antisera over the EC_50_ of mock-depleted antisera for each immunization group, calculated from the ELISA curves in **d**. **f**,**g** Neutralization of either wild-type SARS-CoV-2 (D614G) (**f**) or B.1.351 (Beta) (**g**) pseudotyped lentivirus by mock-depleted, RBD-depleted, or R-KO4-depleted serum. **h**,**i** Normalized NT_50_ for mock-depleted, RBD-depleted, or R-KO4-depleted antisera of each immunization group against SARS-CoV-2 (D614G) (**h**) or B.1.351 (Beta) (**g**) based on the data in **f**,**g**.

To further probe the IgG composition of the polyclonal antisera, we developed a serum-depletion assay in which we could measure the binding and neutralizing titers of antisera selectively depleted from all RBD-binding antibodies except those targeting the S2X259-binding site. The first step involved engineering an RBD protein to bind antibodies from classes 1-3 but not from class 4 (**Fig. 4b**). This protein, R-KO4 (knock-out of class 4 binding), was created by introducing substitutions in nine key positions in the S2X259 epitope. The substitutions were chosen based on examining again the deep mutational-scanning library of the RBD^57^ and predicting which sites in the S2X259 epitope could accommodate amino acids of altered charge (e.g., A372E, T376H, K378N, S383E, R408A, G502D, G504D) or size (e.g., Y369A, T385Y). As the BLI traces show, there is little difference in the binding of class 1,2,3 antibodies between wild-type RBD and R-KO4, whereas binding of five different class 4 antibodies is greatly diminished in R-KO4 (**Fig. 4b**). These results suggest that R-KO4 could be used to deplete sera of class 1,2,3 antibodies and allow us to confirm that our immunofocusing approach biases the immune response to the class 4 epitope.

For our depletion assay, we pooled sera from all five mice in each immunization group. Each vial of pooled antisera (RWT, R11K, R11K-PMD, and R11K-CL) was divided into three samples and mixed with either unconjugated beads (mock-depleted serum), beads bound to RBD (RBD-depleted serum), or beads bound to R-KO4 (R-KO4-depleted serum) (**Fig. 4c**). After allowing IgG in the sera to bind to the beads for 30 min, the beads and bound antibodies were filtered out using spin-filtration columns. Next, using an ELISA assay, we measured binding of each treatment group to RBD (**Fig. 4d**). As expected, mock-depleted antisera from each immunization group bound well to RBD while RBD-depleted antisera showed no appreciable binding in all groups. In the case of R-KO4-depleted antisera, the binding titers varied greatly between the immunization groups. Normalized to the binding titer of mock-depleted serum, the binding titers for R-KO4-depleted antisera were 1% for RWT mice, 6% for R11K mice, and 16% for both R11K-PMD and R11K-CL immunized mice antisera (**Fig. 4e**). These results suggest that immunization with cryptic-face targeting antigens successfully elicits a higher percentage of class 4 antibodies although quantitation of the ELISA titers is convoluted by antibody affinity and quantity.

Finally, we wanted to confirm that the S2X259-like antibodies present in the R11K-PMD and R11K-CL antisera were also functional and contributing to the neutralizing activity we had previously observed (**Fig. 3c**). First, we tested the neutralizing activity of mock-depleted pooled serum from each group against wild-type SARS-CoV-2 and Beta SARS-CoV-2 to establish a baseline of neutralizing activity (**Fig. 4f,g**). As expected based on the ELISA results (**Fig. 4d**), the mock-depleted sera from each immunization group potently neutralized both viruses while the RBD-depleted sera could not neutralize either virus. Consistent with previous reports of minimal class 4 antibodies generated by immunization with wild-type RBD^21,23,24^, the R-KO4-depleted sera from RWT or R11K immunized mice had very low neutralizing activity while the R11K-PMD and R11K-CL groups retained robust neutralizing activity (**Fig. 4f,g**). A comparison of the normalized neutralizing titers of each antisera group following antigen depletion shows that R11K-PMD and R11K-CL antisera had the greatest proportion of functional class 4 antibodies (**Fig. 4h,i**). These results provide good evidence that antigens designed to expose the cryptic-face successfully elicit a more biased, epitope-focused immune response.

## DISCUSSION

Developing vaccines that provide broad coverage across viral strains and are robust to viral evolution is a major challenge in the field of vaccinology. Although current vaccines have helped decrease the morbidity and mortality caused by the recent COVID-19 pandemic, the continuous evolution of the SARS-CoV-2 virus and the threat of cross-species transmission of new *Sarbecoviruses* continues to pose a significant public health threat that could be ameliorated by a universal vaccine. Such universal vaccines are also lacking for many highly variant viruses including influenza, HIV-1, and Ebola.

Immunofocusing is a method to bias the immune response towards broadly neutralizing epitopes by diminishing the immune response towards off-target, non-neutralizing, and subtype-specific immunodominant epitopes on a given antigen.^9^ PMD is an attractive immunofocusing method because it is generalizable and only requires an antigen of interest and a mAb that binds to a target epitope.^51^ In the context of SARS-CoV-2, we initially show that the addition of 11 surface-exposed lysines to the RBD of SARS-CoV-2 did not distort the conformation of the S2X259 epitope. Then, using PMD, we created immunogens designed to have antigenicity at only the class 4 antibody-binding site on the cryptic face of the RBD. Our PMD immunogens were able to generate robust binding titers and broad neutralizing activity in a mouse immunization study. Our serum depletion assay confirmed the ability of PMD to specifically guide the B-cell response towards the S2X259 epitope. Specifically, using this assay, we demonstrated that the proportion and neutralizing activity of antibodies targeting the cryptic face of the RBD was higher in mice immunized with PMD antigens compared to the unmodified RBD antigen (R11K), providing further evidence that immunofocusing with PMD can alter the landscape of the polyclonal antibody response. Interestingly, lysines alone may have an effect on protein antigenicity^61^ and may have altered the landscape of elicited antibodies in R11K immunized mice. However, the addition of stabilizing mutations to R11K, which are not present in wild-type RBD or our RWT antigen, may convolute these results as well as the small sample size of immunized mice leading to lack of statistical significance.

Despite this, the results of our serum-depletion experiments are consistent with studies suggesting that the epitope of class 4 antibodies is subdominant to other regions of the RBD.^21–25^ This subdominance has been explained in part by the fact that the cryptic face is only accessible on the spike protein when the RBD is in the “up” conformation.^62^ However, the results of this study indicate that the subdominant nature of the cryptic face cannot be explained solely by accessibility. Understanding the mechanism behind immunodominance in SARS-CoV-2 as well as other viruses like HIV-1 and influenza is an ongoing area of research and has important implications for vaccine design.^63–66^

From a technological standpoint, it is possible to further optimize PMD and combine it with other emerging vaccine methodologies.^9^ In this work, we demonstrated the feasibility of using Bis NHS-esters to cross-link and stabilize PEG-modified antigens. Cross-linking may be particularly useful in preserving conformation-dependent epitopes in unstable proteins. PMD could also be expanded to include different modifying agents or the use of chemistries outside of NHS-esters. In particular, because PEGs lower the immunogenicity of an antigen^67^, multimerization of PMD antigens onto nanoparticles to boost neutralization titers, or in combination with other immunofocusing techniques like hyperglycosylation and antigen reorientation, could lead to improved vaccine candidates.^68–70^ Additionally, recent results suggest that the prevalence of a single RBD antibody class in polyclonal sera increases the possibility of neutralization escape by viral variants with only a small number of amino acid substitions.^25^ Future efforts to create a pan-*Sarbecovirus* vaccine may benefit from immunofocusing to two synergistic/complimentary epitopes to minimize vulnerability to escape variants at a single site.

The results presented here demonstrate the ability of PMD to focus the antibody response towards a sub-dominant but neutralizing epitope on the RBD of SARS-CoV-2 spike protein. Our serum-depletion assay using an epitope-knock out antigen is also generalizable and could facilitate the evaluation of future immunofocused vaccines. The combination of immunofocusing with other modern vaccine technologies will hopefully aid in the creation of new broadly neutralizing vaccines against not only coronaviruses but also historically intractable viruses like HIV-1 and influenza.

## MATERIALS AND METHODS

### Cell lines

HEK-293T cells were purchased from American type culture collection (ATCC) and maintained in D10 media comprised of Dulbecco’s Modified Eagle Medium (DMEM, Cytiva) supplemented with 10% fetal bovine serum (GeminiBio) and 1% *L*-glutamine/penicillin/streptomycin (GeminiBio). HeLa-ACE2/TMPRSS2 cells were a generous gift from Dr. Jesse Bloom at the Fred Hutchinson Cancer Research Center and were maintained in D10 media. Expi-293F cells (ThermoFisher) were cultured in polycarbonate shaking flasks (Triforest Labware) using a 2:1 v/v mixture of Freestyle293 media and Expi-293 media (ThermoFisher).

### Antibody and antigen cloning

Antibody sequences and Fc-tagged ACE2 were cloned into the CMV/R plasmid backbone for expression under a CMV promoter as previously described.^71^ DNA fragments encoding the variable heavy chain (HC) and light chain (LC) of each antibody (obtained from crystal structures in the RCSB Protein Databank) were codon-optimized and synthesized by Integrated DNA Technologies (IDT). The DNA fragments for each were designed to include a 15 base-pair overlap with the open CMV/R vector. These fragments were then cloned into the CMV/R plasmid containing the VRC01 HC and LC constant domains by In-Fusion (Takara Bio). DNA encoding the wild-type SARS-CoV-2 Wuhan Hu-1 receptor-binding domain (residues 319-533, GenBank MN908947.3) was cloned into an in-house mammalian protein expression vector (pADD2). A hexa-histidine tag and an Avi-Tag^72^ were included on the C-terminus to enable purification and biotinylation. RBD variants encoding either the repackaging substitutions and/or lysine substitutions were cloned by PCR site-directed mutagenesis using wild-type RBD as the backbone. All plasmids were sequence confirmed using Sanger sequencing (Sequetech). To generate DNA for transfection, plasmids were transformed into Stellar^TM^ cells (Takara Bio) and isolated using Maxiprep kits (NucleoBon Xtra Maxi kit, Macherey Nagel). Protein sequences for all antibody variable regions and RBD antigens can be found in the supplemental information. Plasmids for the production of the biotinylated RBDs from WIV-1, Rf1, Yun11, BtKY72, and BM4831 were kindly provided by the Bjorkman lab.

### Protein expression and purification

All proteins were expressed in Expi-293F cells maintained at 37 °C with constant shaking (120 rpm) in a humidified CO_2_ (8%) incubator. Expi-203F cells at a density of 3-4 × 10^6^ cells/mL were transfected using FectoPro transfection reagent (Polyplos) according to the manufacturer’s specifications. Briefly, for a 200 mL transfection, 120 µg plasmid DNA was added to 20 mL media (2:1 v/v mixture of Freestyle293 media and Expi-293 media) and vortexed after the addition of 260 µL FectoPro. The transfection mixture was allowed to incubate at room temperature for 10 min before addition to the Expi-293F cells. Immediately following transfection, cells were boosted with *D*-glucose (4 g/L, Sigma-Aldrich) and valproic acid (3 mM, Acros Organics). The DNA/FectoPro amount was scaled proportionally depending on the size of the transfection. For antibodies, the total DNA amount was the same but was compromised of a 1:1 mixture of heavy-chain plasmid and light-chain plasmid. Biotinylated proteins were produced by transfecting Expi-293F cells with the addition of the BirA enzyme. The transfections was harvested on day 4-5 by centrifuging the cells at 7000 *g* for 5 min. The resulting supernatant was filtered through a 0.22 µm membrane before further purification. For His-tagged proteins, HisPur^TM^ Ni-NTA resin (ThermoFisher Scientific, Cat#88221) was added to filtered supernatant (4 mL/500 mL transfection) and 1M imidazole in PBS (10 mM phosphate, 2.7 mM KCl, pH 7.4, Bioland Scientific LLC, Cat# PBS01-03) was added to the supernatant to a final concentration of 5 mM imidazole to minimize binding of impurities. After allowing the resin to bind overnight at 4 °C with gentle spinning, the mixture was passed over a gravity-flow column. The collected resin was washed 3 x 10 mL with 20 mM imidazole in PBS before eluting with 15 mL of 250 mM imidazole in PBS. The elution was concentrated using Amicon^R^ centrifugal filters (10 kDa MWCO for RBD, Millipore Sigma) and then purified on an ÄKTA pure^TM^ chromatography system (Cytiva) with a Superdex 200 Increase size-exclusion chromatography column (Cytiva) in PBS. Peak fractions were collected based on the chromatography trace, measuring absorbance at 280 nm. All antibodies (and ACE2-Fc) were purified by first diluting the filtered Expi-293F supernatant 1:1 with 1 × PBS before direction application onto a 5mL HiTrap^TM^ MabSelect^TM^ PrismA column (Cytiva) using an ÄKTA pure^TM^ chromatography system (Cytiva). Post protein binding, the column was washed with PBS and then protein was eluted with 15 mL 100 mM glycine in PBS (pH 2.8) into 1.5 mL 1M Tris pH 8.0. The eluent was then concentrated and buffer exchanged into PBS. The concentration of all proteins was determined by absorbance at 280 nm, and purity and size confirmed by protein gel electrophoresis. Protein samples were flash-frozen in PBS with 10 % glycerol before for storage at −20 °C or −80 °C.

### Screening for lysine expression

Based on the structure the SARS-CoV-2 RBD (PDB ID: 7M7W) as well as the RBD deep-mutational scanning data,^57^ we initially identified 11 surface-exposed sites to introduce a lysine into via site-directed mutagenesis. The expression levels of the single-substituted RBD variants in Expi-293F cells were compared via immunoblotting. Three days after transient transfection, Expi-293F cultures were centrifuged at 7,000 *g* for 5 min before supernatant samples were collected. Supernatants from each culture (5 µL) were directly pipetted onto 0.2-µm nitrocellulose membranes (Bio-Rad) and allowed to dry for 20 min. The membrane was then blocked for 20 min using PBST (10 mM phosphate, 2.7 mM KCl, 0.05% Tween-20, pH 7.4) with milk (10% *w/v* non-fat dry milk, Bio-Rad) before S2X259 antibody (1:2500 in PBST with milk) was added for a 1 h incubation. Following primary antibody, the blot was rinsed with 3 x 20 mL PBST before adding HRP-conjugated rabbit anti-human IgG (1:5000 dilution in PBST with milk) for another 1 h incubation. Following another 3 x 20 mL wash with PBST, the blot was developed using a Western blotting substrate (Pierce ECL, Thermo Scientific) and read on a GE Amersham Imager 600. The resulting image was analyzed with Fiji (ImageJ v.2.1.0) and expression level of RBD variants was compared to the expression of wild-type RBD. A second round of lysine screening was conducted in the same way, except that additional single lysine substitutions were made to an RBD protein already containing 7 additional lysines (RBD+7K) and expression level was compared to the expression level of RBD+7K.

### Differential scanning fluorimetry

Protein thermal-melting profiles were obtained using a Prometheus NT.48 Instrument (NanoTemper). Proteins were diluted in PBS to a concentration of 0.1 mg/mL and loaded into glass capillaries (NanoTemper). Samples were then heated at a rate of 1 °C per min in a gradient that ranged from 20 to 95 °C while intrinsic fluorescence was recorded (350 nm and 330 nm). The thermal melting curve was obtained by plotting the first derivative of the ratio (350 nm/330 nm). The melting temperature was calculated automatically by the instrument (PR.ThermControl software). Thermal melting profile for each protein was obtained in triplicate.

### S2X259 Fab generation

To generate the fragment-antigen binding (Fab) of S2X259, 50 µL of 1M Tris pH 8.0 was added to 0.5 mL of S2X259 IgG at 2 mg/mL in PBS. To this mixture, 2 µL of Lys-C was added for each mg of IgG and allowed to digest for 1.5 h at 37 °C with gentle spinning. Digestion was terminated by the addition of 25 µL of 10% of acetic acid. To remove the Fc portion of the IgG as well as any undigested IgG, protein G (ThermoFisher Scientific, Cat# 20397) was added to the mixture and allowed to incubate overnight at 4 °C with gentle spinning. The next day, the protein G resin was removed by gravity filtration over a column. The flow-through was collected, equilibrated with 10 × PBS and then purified on an ÄKTA pure^TM^ chromatography system (Cytiva) with a Superdex 200 Increase size-exclusion chromatography column (Cytiva). Size and digestion of the purified Fab was confirmed by gel electrophoresis.

### Biolayer interferometry (BLI)

Biolayer interferometry was performed on an Octet^TM^ RED96e system (Pall FortéBio) in 96-well flat-bottom black plates (Greiner). All samples were run in PBS with 0.1% BSA and 0.05% Tween-20 and assays were performed under agitation (1000 rpm). Antibodies were loaded onto tips using the anti-human IgG Fc capture (AHC) Biosensors (FortéBio), biotinylated antigens were loaded onto Streptavidin Biosensors (Sartorius), and His-tagged proteins were loaded onto NTA Biosensors (FortéBio). After loading, tips were dipped into wells containing buffer only for 20 sec, before dipping into wells containing a binding partner (association) followed by dipping into wells containing buffer (dissociation). All samples in all experiments were baseline-subtracted to a well that contained an antigen-loaded tip that was dipped into a well without a binding partner to control for any buffer trends between the samples. The baseline subtracted binding curves (processed by FortéBio Data Analysis software) were exported and plotted in GraphPad Prism.

### S2X259 affinity resin column coupling

To produce the S2X259 affinity resin column, AminoLink^TM^ Plus coupling resin (ThermoFisher Scientific, Cat# 201501) with the pH 7.2 coupling protocol was used. Briefly, 10 mg of S2X259 antibody in 3 mL PBS was added to 1 mL of AminoLink^TM^ Plus coupling resin pre-equilibrated in PBS. To this mixture, 40 µL of 5M NaCNBH_3_ in 1M NaOH was added and the mixture was left to react at 4 °C overnight with gentle rotation. The following day, the resin was filtered and the protein concentration of the flow-through was measured to confirm coupling. The resin was then washed with 10 mL 1 M Tris pH 8.0 and then remaining reactive sites were quenched by incubation of the resin with 3 mL 1 M Tris pH 8.0 and 40 µL of NaCNBH_3_ for 30 min at room temperature. The resin was then washed with 3 × 10 mL 1M NaCl to remove unconjugated protein and then 3 × 10 mL PBS to re-equilibrate the column. S2X259-affinity resin was used immediately in the protect, modify, deprotect (PMD) protocol and was never stored or re-used.

### Protect, modify, deprotect (PMD)

R11K protein in PBS was added to S2X259 affinity resin column and allowed to bind for 10 min at room temperature with gentle spinning. The added amount of R11K protein was calculated based on the estimated amount of bound S2X259 antibody in each column and a theoretical binding of two RBDs per IgG. Binding was confirmed by measuring protein content in the flow-through. The column was washed with 10 mL PBS and then 5 mL of PBS containing a PEG-NHS ester reagent was added such that there was at least 5 × molar excess of PEG moieties per theoretical exposed (free) lysine residue. This was incubated for 20 min at room temperature with gentle rotation, the column drained, and then the incubation repeated with another added equivalent of PBS and NHS-PEG reagent. Following the second incubation, the reaction mixture was eluted, and the column washed with 2 × 10 mL 100 mM Tris pH 8.0 to quench unreacted NHS esters and wash out hydrolyzed NHS esters. The modified RBD protein was eluted with 3 × 3 mL 100 mM glycine pH 1.5 directly into 3 mL of 1M Tris pH 8.5. This solution containing the RBD protein was immediately concentrated, buffer-exchanged, and purified on an ÄKTA pure^TM^ chromatography system (Cytiva) with a Superdex 200 Increase size-exclusion chromatography column (Cytiva). To produce cross-link modified R11K protein, S2X259 affinity resin was first reacted with N-acetyl succinimide to acetylate exposed lysines on S2X259. Briefly, N-acetyl succinimide in 5 mL PBS was added to the resin column such that the acetyl moieties would be in >10 × molar excess of the theoretical calculated lysines of all column-bound S2X259 given an approximation of 75 lysines per IgG. The reaction was allowed to proceed for 30 min to 1 h at room-temperature with gentle rotation. The reaction mixture was then removed and the column quenched with 2 × 10 mL 100 mM Tris pH 8.0 followed by a re-equilibration with PBS. The subsequent steps involving R11K binding and protection, modification, and deprotection were the same as described above other than the usage of a Bis-NHS ester to cross-link exposed lysines on R11K. All PEG-NHS ester reagents were purchased from Quanta Biodesign Ltd: Bis-dPEG^TM^ -NHS ester (Cat# 10224), m-dPEG^TM^ -NHS ester (Cat# 10211).

### SDS-PAGE

Protein samples were prepared by mixing with 4× Laemmli sample buffer (BioRad Cat# 1610747) before boiling at 95 °C for 5 min. 10 µL of each protein sample was then loaded and run on precast protein gels (4-20% Mini-PROTEAN® TGX^TX^, BioRad) before staining with Coomassie blue dye. For a reducing gel, protein samples were mixed with 3× sample buffer (3 parts 4× Laemmli sample buffer mixed with 1 part ß-mercaptoethanol).

### Mouse immunization studies

All mice were maintained in accordance with the Public Health Service Policy for “Human Care and Use of Laboratory Animals” under a protocol approved by the Stanford University Administrative Panel on Laboratory Animal Care (APLAC-33709). Immunizations were performed in female BALB/c mice (6-8 weeks) (*n=*5 per group) *via* intramuscular injection. Each injection was 100 µL containing 5 µg protein antigen adjuvanted with 500 µg Alum (Alhydrogel, InvivoGen) and 20 µg CpG (ODN1826, InvivoGen) and volume adjusted with Dulbecco’s phosphate-buffered saline (Gibco). Mice were immunized on day 0 and boosted on day 35, day 70, and day 105. Mice were bled on day 77 (1 week post boost 2) and day 112 (1 week post boost 3). Blood samples were collected into serum gel tubes (Sarstedt), centrifuged at 10,000 *g* for 15 min, and collected sera were stored at −80 °C.

### Serum enzyme-linked immunosorbent assay (ELISA)

Nunc MaxiSorp 96-well plates were coated with streptavidin at 5 µg/mL in 50 mM bicarbonate pH 8.75 for 1 h at room temperature before washing 3 times with 300 µL of MilliQ-H_2_O using a plate washer (ELx405 BioTek). Plate wells were then blocked with ChonBlock Blocking/Dilution ELISA Buffer (Chondrex) at 4 °C overnight (120 µL per well). Chonblock was then removed before the addition to each well of 50 µL biotinylated-antigen at a concentration of 2 µg/mL. Dilution buffer for antigen and all subsequent reagents was PBST. All subsequent washes were done with PBST (300 µL, 3 times) unless otherwise noted and all incubations were performed at room temperature. After incubation with biotinylated antigen for 1 hour, plates were washed and then 50 µL of serially diluted mouse antisera was added to the 96-well plates and incubated for 1 hour. Antisera was diluted to a starting concentration of 1:50 followed by serial 5-fold dilutions. Following plate washing, 50 µL goat anti-mouse HRP-conjugated IgG (1:5,000) was added and incubated for one hour. Plates were then washed 6 times (300 µL each time) and developed using 50 µL One-Step Turbo TMB substrate (Pierce) for 5 min before quenching with 50 µL 2 M sulfuric acid. Absorbance at 450 nm (A450) was measured using a Synergy BioHTX plate reader (BioTek). For the competition ELISAs, competing antibodies (either S309 or S2X259 at 200 nM) were added to streptavidin coated plates bound with biotinylated-RBD and incubated for 1 hour at room-temperature before washing and addition of serially diluted mouse antisera. For the antibody avidity ELISAs, the same protocol was followed as described above except that following incubation with serially diluted mouse antisera, 2M NaSCN was added to the plates for 15 min at room temperature before washing with PBST.

### Production of pseudotyped lentiviruses

Spike-pseudotyped lentiviruses encoding a luciferase-ZsGreen reporter were produced using a five-plasmid system as previously described.^73^ The plasmid system uses the packaging vector pHAGE-Luc2-IRES-ZsGreen(NR-52516), three helper plasmids HDM-Hgpm2 (NR-52517), HDM-tat1b (NR-52518), pRC-CMV-Rev1b (NR-52519), and a plasmid encoding the viral spike protein. The wild-type SARS-CoV-2 spike plasmid (HDM-SARS2-spike-delta21, Addgene #155130) was cloned with the additional D614G substitution. B.1.351 (Beta) spike included L18F, D80A, D215G, Δ242-244, R246I, K417N, E484K, N501Y, D614G, A701V, and P1 (Gamma) included L18F, T20N, P26S, D138Y, R190S, K417T, E484K, N501Y, D614G, H655Y, T1027I. One day prior to transfection, HEK-293T cells were seeded in 10 cm Petri dishes at a density of 5 × 10^6^ cells. The following day, the plasmids were added to 1 mL of D10 media (10 µg packaging vector, 3.4 µg spike protein, 2.2 µg of each helper plasmid) after which 30 µL of BioT (Bioland Scientific) was added to form transfection complexes. Transfection mixture was incubated for 10 min at room temperature and then added to the HEK-293T cells in their culture dish. After 24 hours culture medium was replenished and 72 h post transfection the culture medium was collected, filtered through a 0.45-µm filter, aliquoted, flash frozen in liquid nitrogen, and then stored at −80 °C.

The non-SARS-CoV-2 *Sarbecovirus* spike sequences were obtained from GenBank and cloned into the spike protein plasmid: SARS-CoV-1 (GenBank: ABD72984.1), pangolin coronavirus PCoV_GD (GenBank: QLR06867.1), WIV-1 (GenBank: AGZ48828.1). These lentiviruses were produced using the same method described above but using Expi-293F cells. One day prior to transfection, Expi-293F cells were diluted to 3 × 10^6^ cells/mL in 200 mL. Transfection mixture was prepared by adding 200 µg packaging vector, 68 µg spike plasmid, and 44 µg helper plasmids to 20 mL FreeStyle293/Expi-293 media followed by the dropwise addition and mixing of 600 µL BioT. After 10 min incubation at room temperature the transfection mixture was added to the Expi-293F cells after which cells were boosted with *D*-glucose (4 g/L) and valproic acid (3 mM). After 72 h the cells were spun down, culture supernatant was collected, filtered through a 0.45-µm filter and then aliquoted, flash frozen in liquid nitrogen, and stored at −80 °C. All pseudotyped lentiviruses were titrated in HEK-293T cells prior to usage.

### Neutralization assays using pseudotyped lentivirus

HeLa cells stably overexpressing human angiotensin-converting enzyme 2 (ACE2) as well as transmembrane serine protease 2 (TMPRSS2) were seeded at a density of 1 × 10^4^ cells/well in white-walled 96-well plates (ThermoFisher Scientific) one day prior to usage (day 0). On day 1, heat-inactivated mouse antisera (56 °C, 30 min) was serially diluted in D10 media and mixed with pseudotyped lentivirus diluted in D10 media supplemented with polybrene (Sigma-Aldrich) at a final concentration of 5 µg/mL). After 1 h incubation at 37 °C the antisera/viral dilutions were transferred to the HeLa/ACE2/TMPRSS2 cells. On day 3, the medium was removed from the wells, and cells were lysed by the addition of 100 µL of luciferase substrates (BriteLite Plus, Perkin Elmer). Luminescent signals were recorded on a microplate reader (BioTek Synergy^TM^ HT or Tecan M200) after shaking for 30 s. Infection percent was normalized to the signal in cells only wells (0% infection) and virus only wells (100% infection) on each plate. The neutralization titer (NT_50_) was defined as the serum dilution concentration at which 50% virus infection was measured. Neutralization assays were performed in technical duplicates.

### Conjugation of biotinylated-RBD to agarose beads

High capacity NeutrAvidin^TM^ beaded agarose (Thermo Scientific, Cat# 29204) were rinsed 3 × in PBS followed by incubation in PBS with 5 % bovine serum albumin (BSA) for 20 min up to 1 h. Beads were rinsed 3 × in PBS before addition of biotinylated-protein. Biotinylated-protein was added in an amount equal to 75% of the theoretical maximum binding capacity of the agarose, according to the manufacturer’s specifications. Following incubation at room temperature for 30 min with gentle rotation, agarose beads were washed 4 × in PBS to remove unbound protein. Agarose beads were used immediately and created fresh for every usage.

### Serum antibody depletion experiment

Heat-inactivated mouse antisera (56 °C, 30 min) was pooled from mice in each immunization group (*n*=5 mice per group) and diluted with D10 media to a concentration of 1:50 (ex. 3 µL antisera from each mouse added to 735 µL of media). Pooled serum was then divided evenly into three tubes and the same volume of agarose beads was added to each tube. Washed agarose beads were added to one tube to create the mock-depleted sample, RBD-bound agarose beads were added to the second tube, and R-KO4-bound agarose beads were added to the third tube. The volume of agarose beads added was normalized to correspond to approximately 0.2-0.3 mg of RBD protein each time. Beads and antisera were incubated for 30 min at room temperature with gentle rotation before the mixtures were filtered through a 0.22 µm filter to remove the beads. The collected diluted antisera was used directly for neutralization assays or ELISAs.

### Statistical analyses

All normalization, curve-fitting, and statistical analyses were performed using GraphPad Prism 9.5.1 software. Log-transformed data (ELISA titers and NT_50_) are presented as geometric mean ± standard deviation. Comparison between two groups was performed using the two-tailed Mann-Whitney U test. *P* values of 0.05 or less are considered significant and plotted.

### Data Availability

All data supporting the results in this study are available within the main text and its supplementary information.

## Acknowledgments

We thank all members of the Kim lab for helpful discussion and guidance during this project. We thank Dr. Abigail Powell, Dr. Mrinmoy Sanyal, Dr. Natalia Friedland, and Ya-Chen Cheng for working tirelessly during the COVID-19 pandemic to set up the SARS-CoV-2 neutralization assay and produce so many of the SARS-CoV-2 pseudotyped lentiviruses and variants of concern. We also thank the Bjorkman lab for generously sharing their plasmids with us. Thank you to Dr. Duo Xu for constant support and assistance with mouse immunizations at critical moments. Thank you to Dr. Duo Xu, Dr. Soohyun Kim, and Ashley Utz for comments on the manuscript. T.U.J.B was supported by the Knight-Hennessy Graduate Scholarship and a Canadian Institutes of Health Research Doctoral Foreign Study Award (FRN:170770). This work was supported by an NIH Director’s Pioneer Award (DP1-AI158125), the Virginia & D.K. Ludwig Fund for Cancer Research, and the Chan Zuckerberg Biohub to P.S.K.

## Author contributions

Conceptualization, methodology, interpretation: T.U.J.B, P.A.W.B, P.S.K. Experiments: T.U.J.B. Mouse immunization studies: J.D. Writing (original draft): T.U.J.B. Writing (revised draft): T.U.J.B., P.S.K. Writing (review & editing): T.U.J.B, P.A.W.B, P.S.K.

## Supporting Information

**Figure S1.**
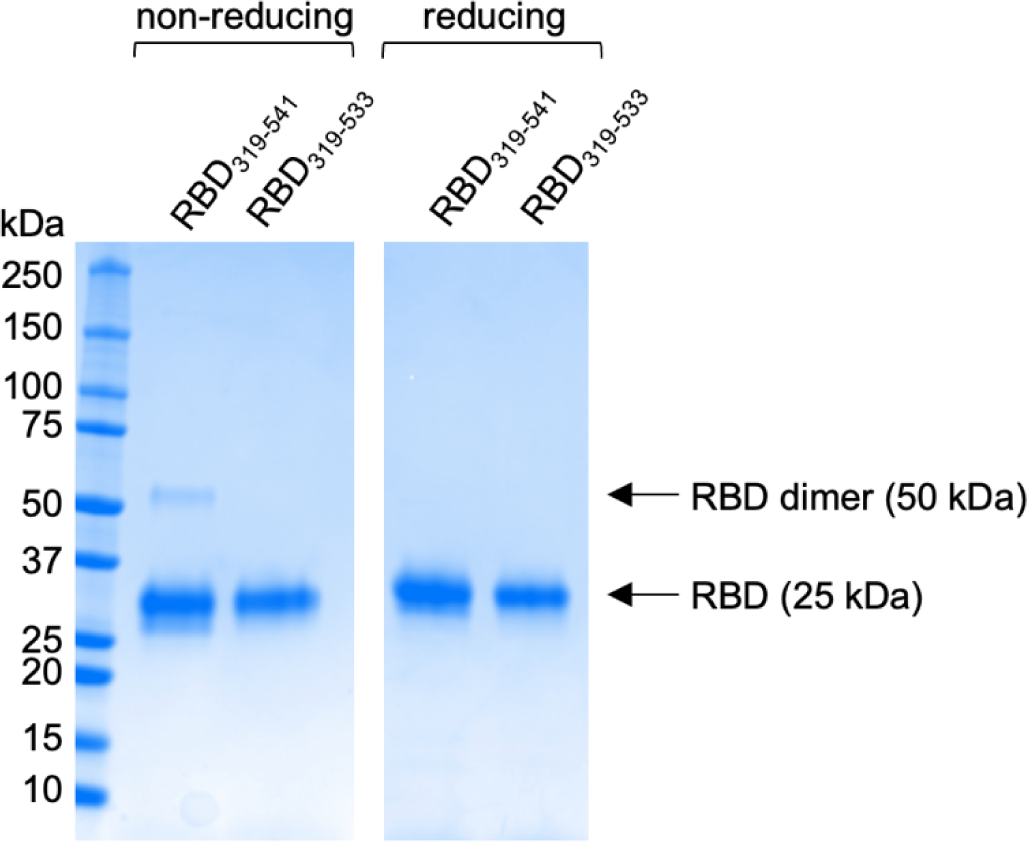
Removing the C-terminal cysteine from RBD substantially reduces dimer formation. Reducing and non-reducing SDS-PAGE gel of the RBD of Wuhan Hu-1 SARS-CoV-2 (GenBank MN908947.3) containing either residues 319-541 or residues 319-533 with one fewer cysteine residue.

**Figure S2.**
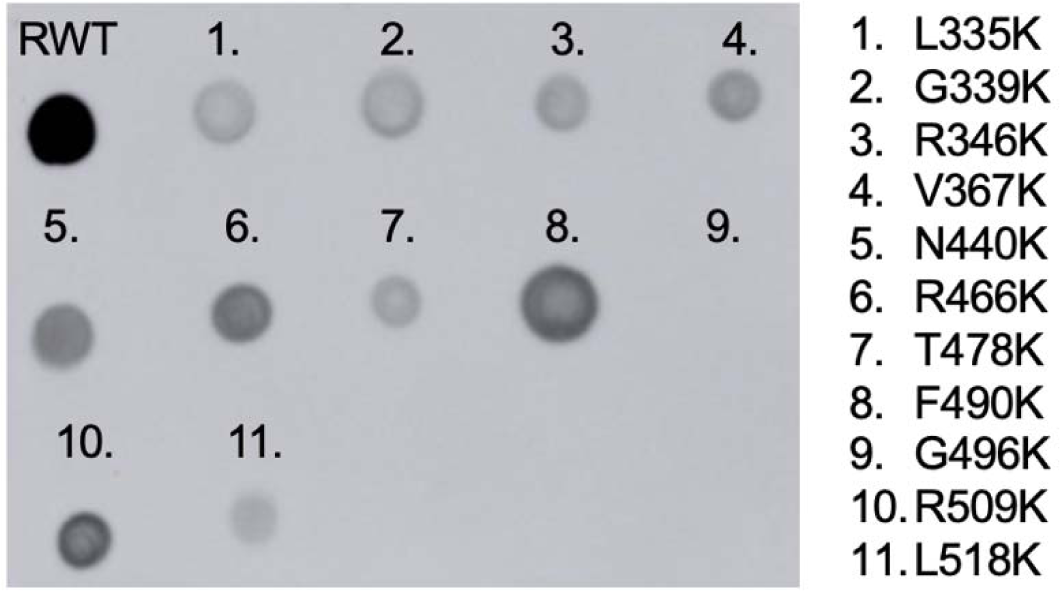
Expression levels of wild-type RBD and RBD variants containing lysine substitutions. A representative dot-blot is shown to compare the expression levels of wild-type RBD (RWT) and RBD variants from small-scale expressions. Supernatants from each cell culture of lysine-substituted RBDs (5 µL) were directly pipetted onto 0.2-µm nitrocellulose membranes before blocking and incubation with mAb S2X259 and detection with HRP-conjugated rabbit anti-human IgG. The blot was imaged on a GE Amersham Imager 600 and analyzed with Fiji (ImageJ v.2.1.0). The 7 lysine substitutions that were chosen to create R7K and their relative expression-level compared to RWT are: #3 R346K (59%), #4 V367K (47%), #5 N440K (163%), #6 R466K (104%), #7 T478K (38%), #8 F490K (142%), #11 L518K (33%).

**Figure S3.**
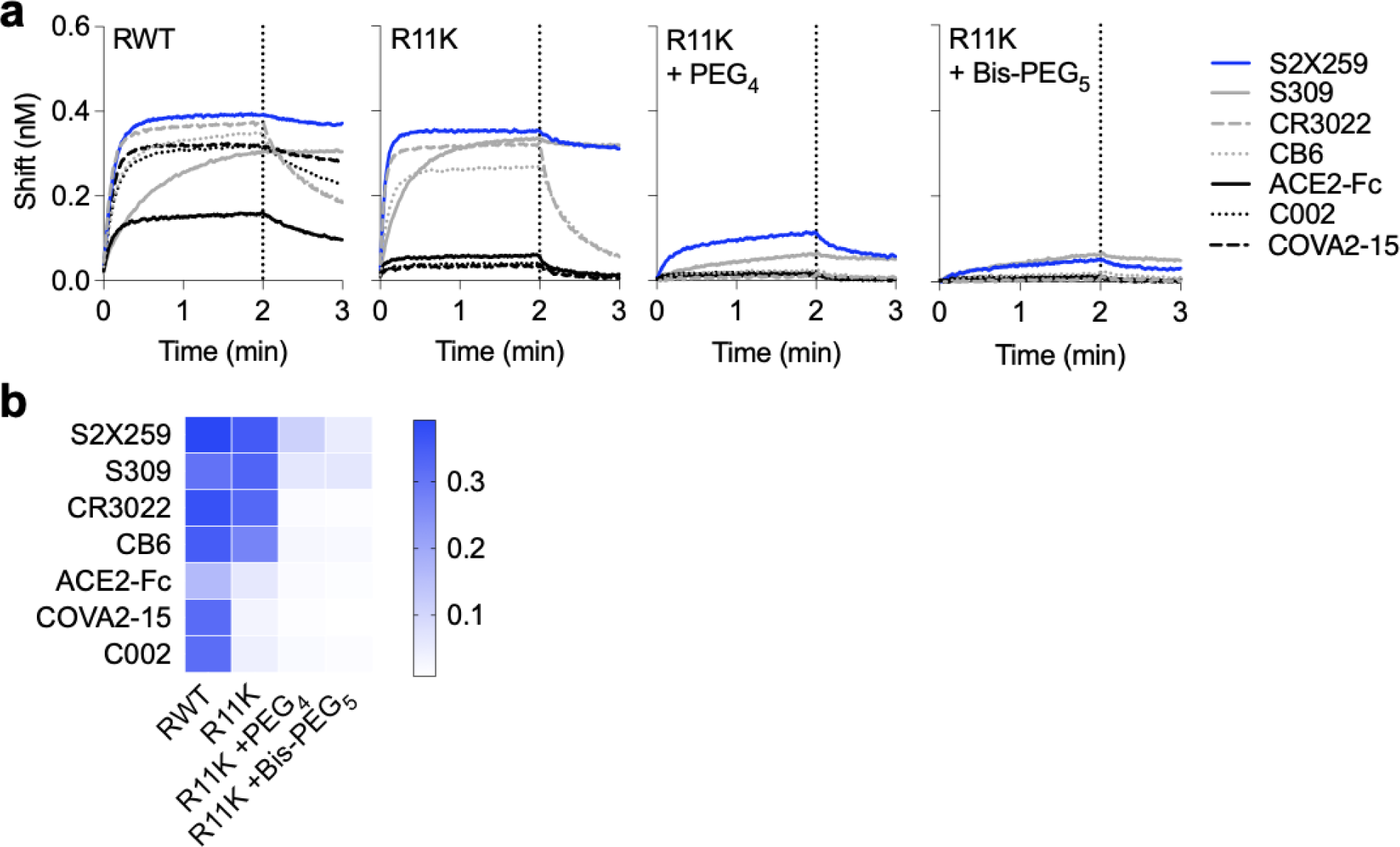
Modification of R11K prior to protection severely decreases binding to S2X259. R11K was reacted with NHS-PEG_4_ or Bis-NHS-PEG_5_ in solution, omitting the PMD protection step in which R11K is bound to an S2X259-conjugated resin. After reaction for 30 min, excess PEG moieties were removed by size-exclusion chromatography. **a**, BLI binding curves of RWT, R11K, and the purified PEGylated proteins (R11K+PEG_4_ and R11K+Bis-PEG_5_) binding to a panel of RBD-directed antibodies, including S2X259, S309 and CR3022. **b**, BLI association amplitude of each antibody binding to RWT, R11K, R11K+ PEG_4_, R11K+Bis-PEG_5,_ as shown in **a**.

**Figure S4.**
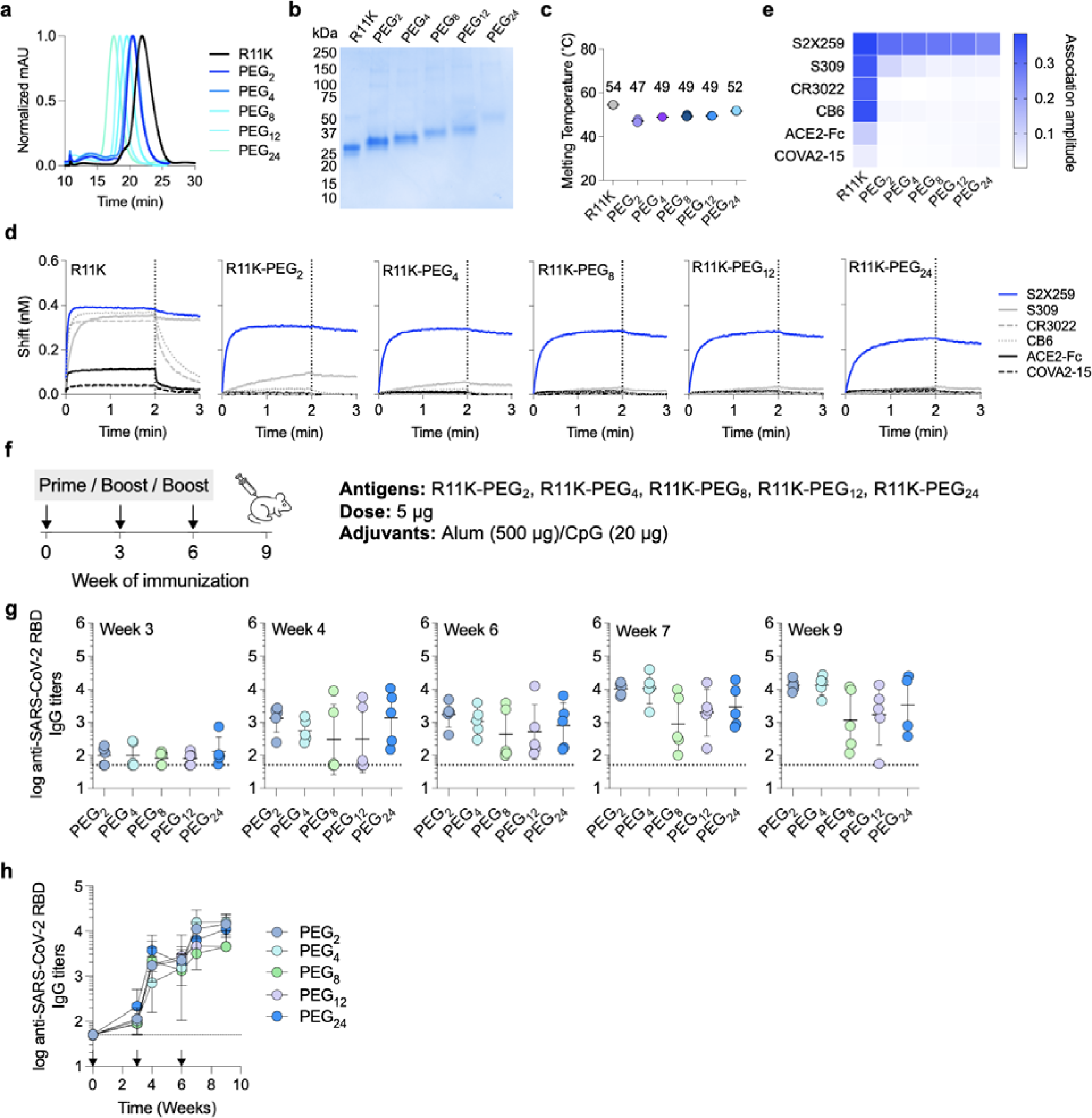
Antigenicity of R11K modified with different PEG lengths. **a**, Normalized size-exclusion chromatographic traces of R11K and R11K variants modified with PEG chains of differing lengths (*n* = 2,4,8,12, or 24). **b**, SDS-PAGE analysis of R11K and PEG-modified R11K variants post-expression and purification via size-exclusion chromatography. **c**, Thermal melting temperature for R11K, R11K-PEG_2_, R11K-PEG_4_, R11K-PEG_8_, R11K-PEG_12_, and R11K-PEG_24_ measured by differential scanning fluorimetry. Data are presented as mean ± standard deviation (*n* = 3 replicates). **d**, Binding curves of RBD-directed antibodies from all four classes to R11K, R11K-PEG_2_, R11K-PEG_4_, R11K-PEG_8_, R11K-PEG_12_, and R11K-PEG_24_. **e**, BLI association amplitude of each antibody binding to the R11K variants. Association was monitored for 2 min (dotted lines), after which dissociation was monitored for 1 min. **f**, Schematic of the mouse immunization with a three-dose regimen occuring on days 0, 21, and 42. Mice were immunized with 5 *ug* of antigen adjuvanted with alum/CpG (500 *u*g/20 *u*g) *via* intramuscular injection (*n* = 5 per group). **g**, Serum IgG titers to SARS-CoV-2 RBD at 3, 4, and 6 weeks post-boost 1 (the latter two corresponding to 1 and 3 weeks post-boost 2). Each circle represents an individual mouse and the mean titers are indicated with a dash. **h**, Serum IgG titers to SARS-COV-2 RBD over time. Horizontal dotted lines indicate the limits of quantification. Data are presented as geometric mean ± standard deviation.

**Figure S5.**
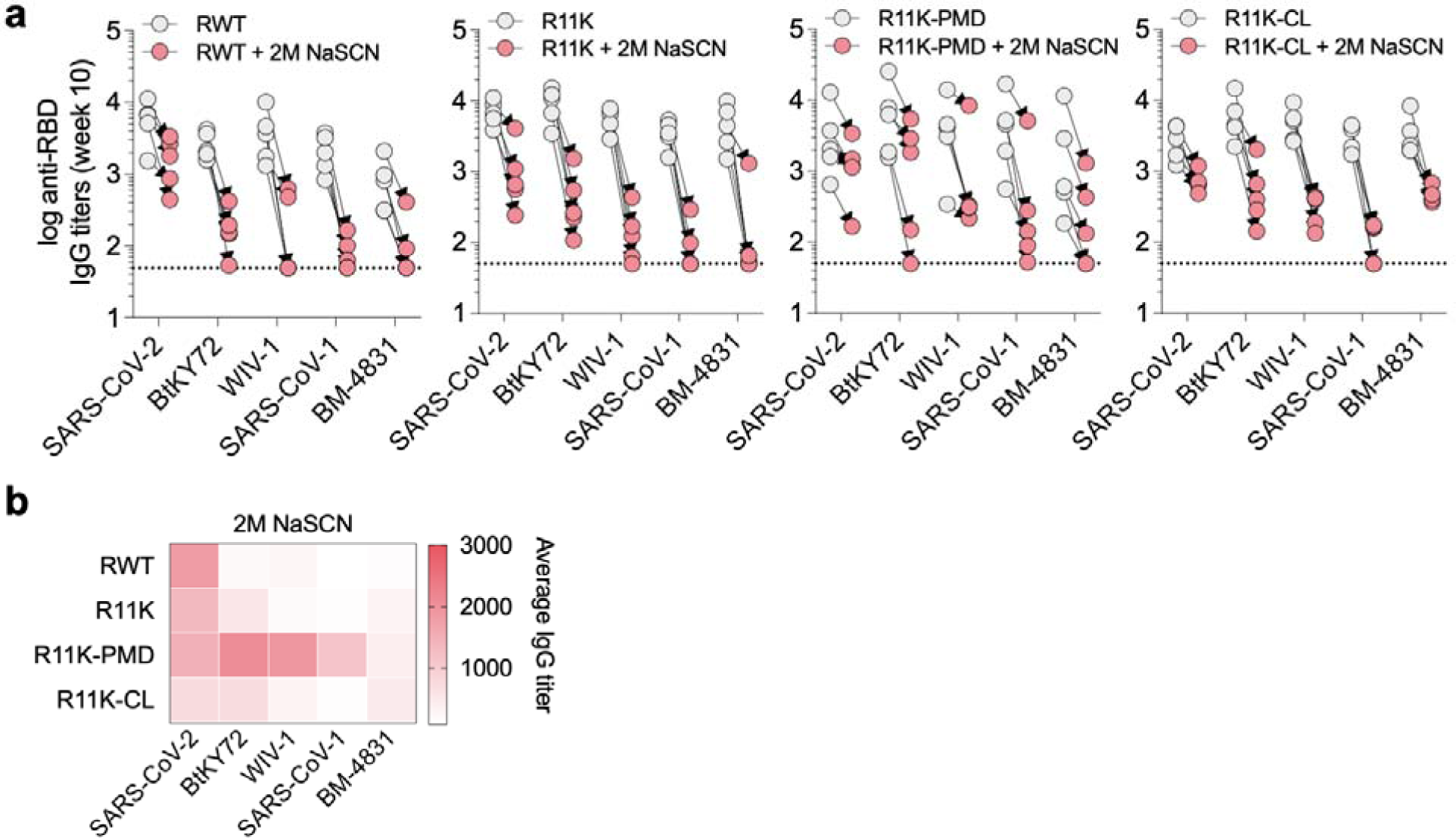
Antibody-avidity assay. **A**, Serum IgG titers against different biotinylated RBDs in the presence or absence of treatment with 2M sodium thiocyanate (NaSCN). Each circle represents a single mouse. Horizontal dotted lines indicate the limit of quantitation. **B**, Average serum IgG titers of each immunization group to different RBDs post-treatment with 2M NaSCN presented as a heat map.

**Figure S6.**
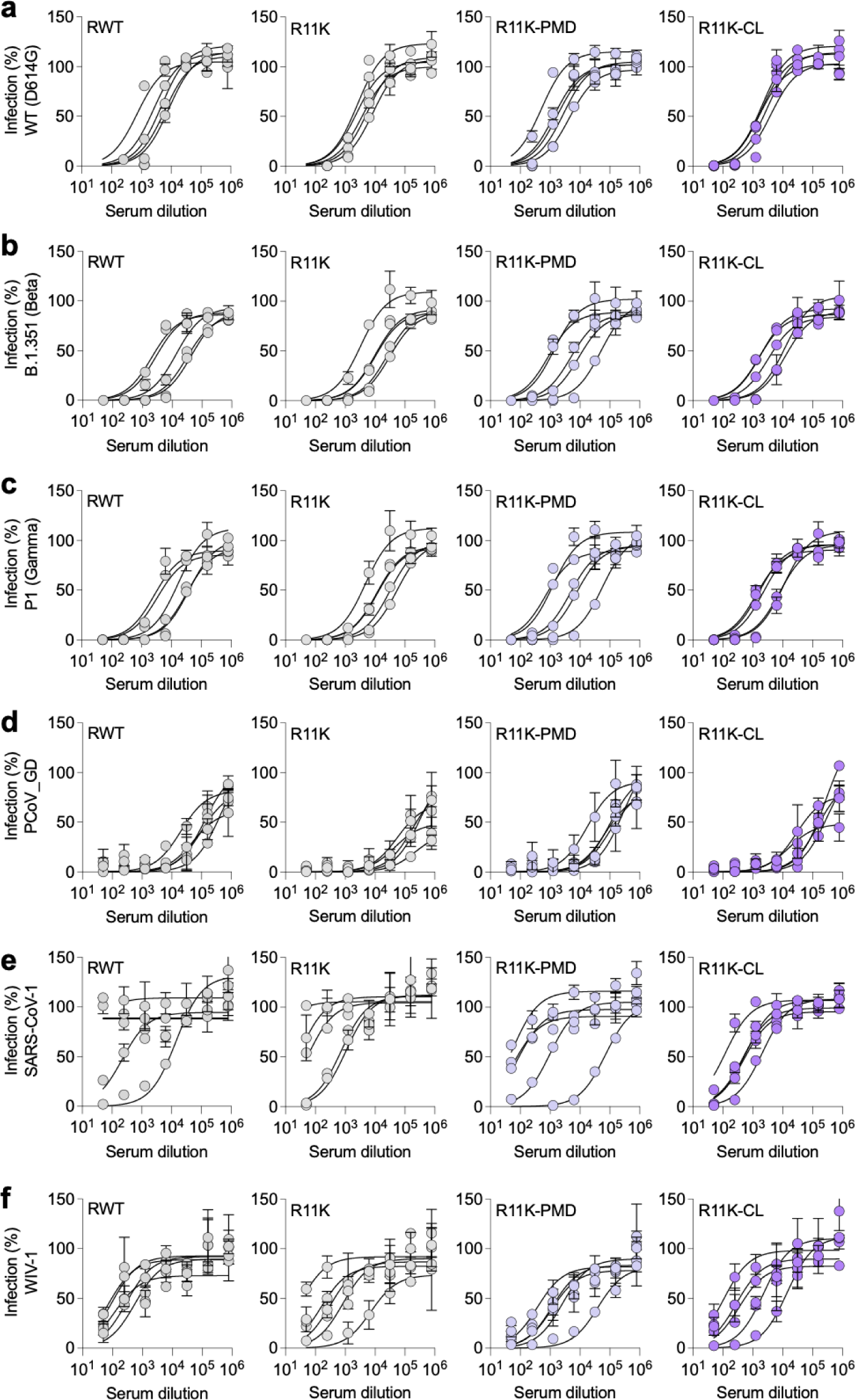
Neutralization of *Sarbecovirus* spike-pseudotyped lentiviruses. Antisera from mice immunized with either RWT, R11K, R11K-PMD, or R11K-CL were assessed on day 112 for neutralizing activity in HeLa cells overexpressing ACE2 and TMPRSS2. Neutralization curves are shown for each immunization group against wild-type SARS-CoV-2 (D614G) (**a**), SARS-CoV-2 Beta (**b**), SARS-CoV-2 Gamma (**c**), pangolin coronavirus PCoV_GD (**d**), SARS-CoV-1 (**e**), and WIV-1 (**f**). Data are presented as mean ± standard deviation (*n*=2 technical duplicates). Each curve represents a single mouse.

### Supplementary Tables

#### Antibody Sequences

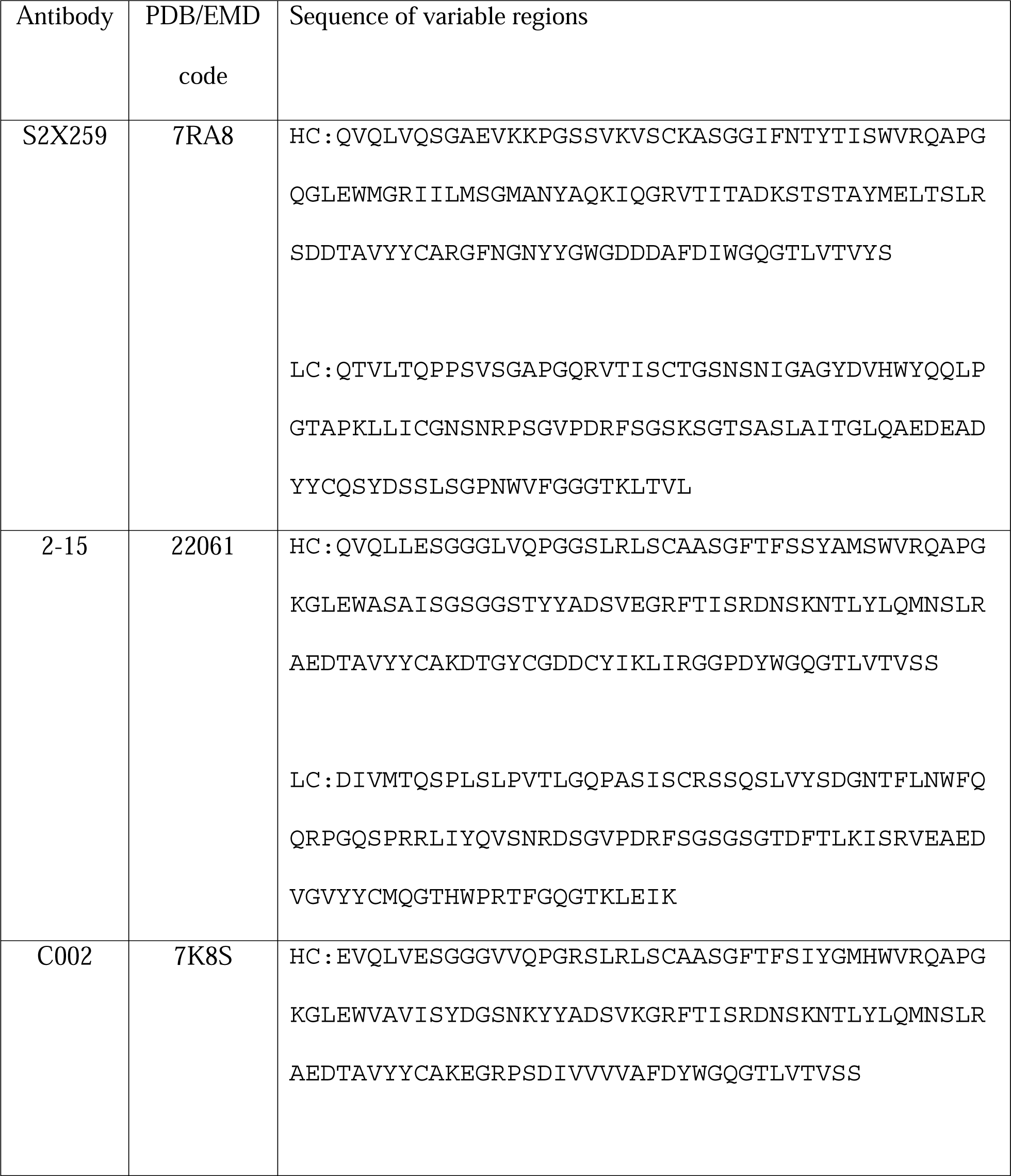
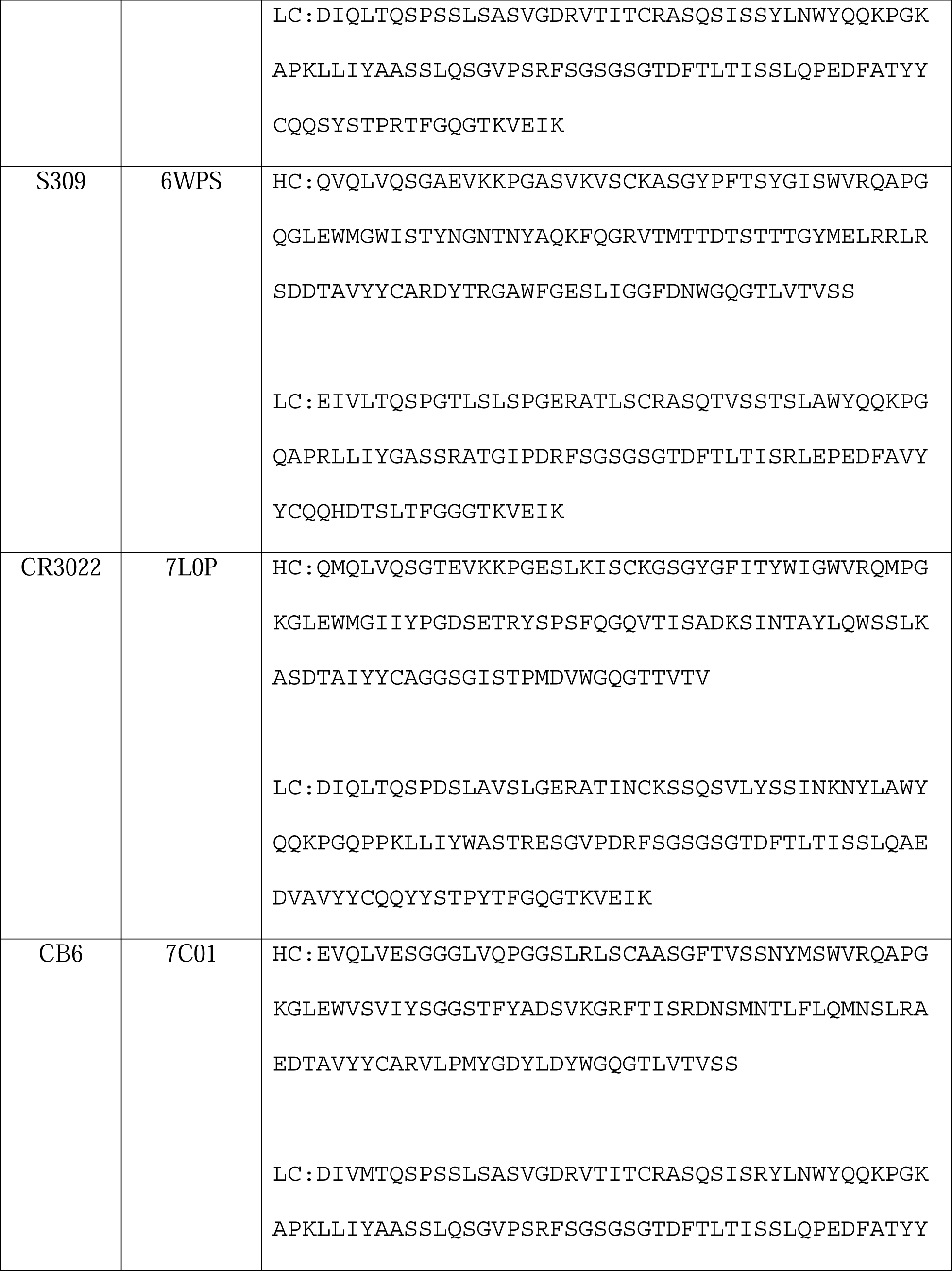
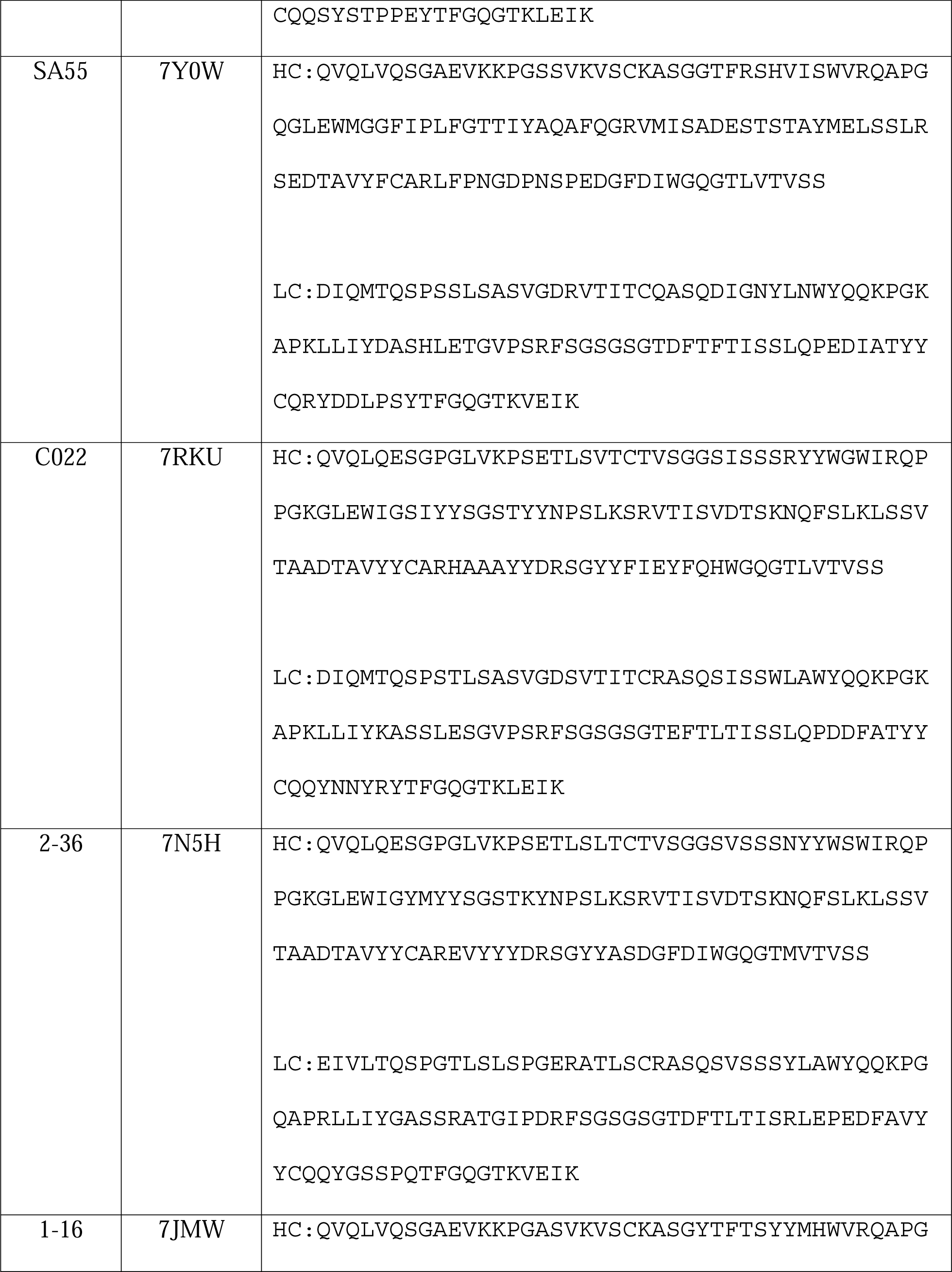
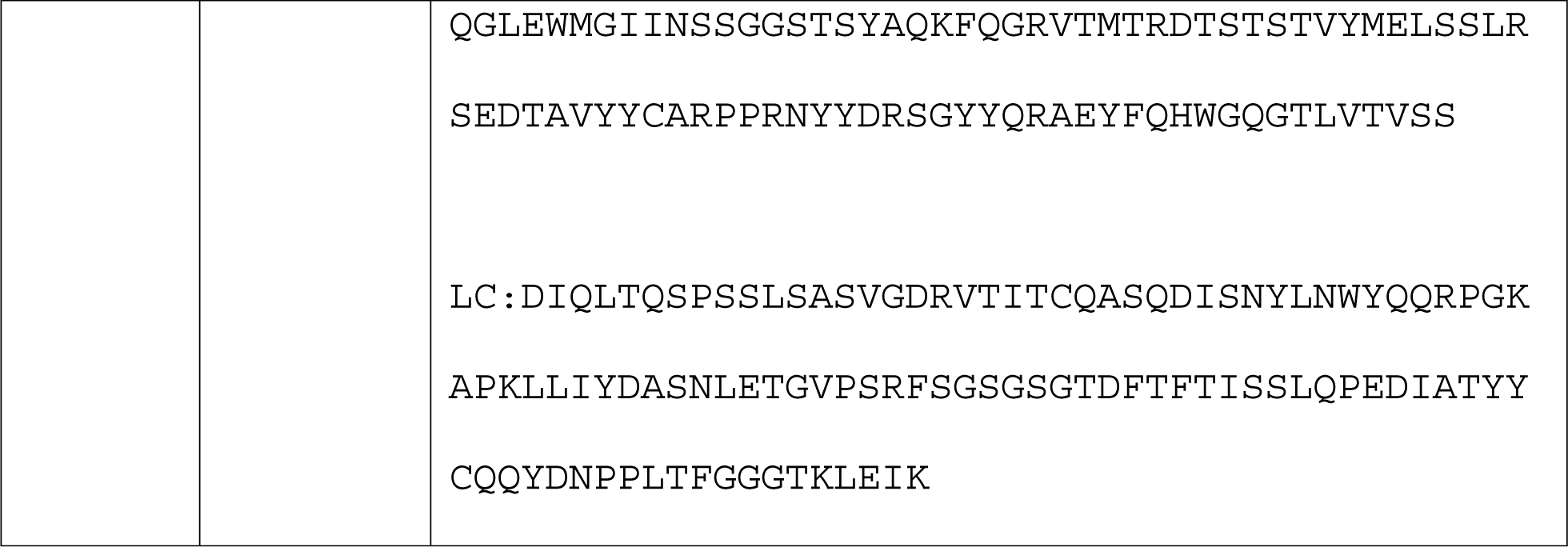

#### RBD Sequences

<u>Hexa-histidine tag</u> (underlined residues)

**Avi-Tag** (boldfaced residues)

*Fc region* (italicized residues)

***TEV site*** (boldface italicized residues)

#### SARS-CoV-2 RBD (RWT)

RVQPTESIVRFPNITNLCPFGEVFNATRFASVYAWNRKRISNCVADYSVLYNSASFSTFKCYGV SPTKLNDLCFTNVYADSFVIRGDEVRQIAPGQTGKIADYNYKLPDDFTGCVIAWNSNNLDSKVG GNYNYLYRLFRKSNLKPFERDISTEIYQAGSTPCNGVEGFNCYFPLQSYGFQPTNGVGYQPYRV VVLSFELLHAPATVCGPKKSTNLG<u>HHHHHH</u>G**GLNDIFEAQKIEWHE**

#### R7K

RVQPTESIVRFPNITNLCPFGEVFNATKFASVYAWNRKRISNCVADYSKLYNSASFSTFKCYGV SPTKLNDLCFTNVYADSFVIRGDEVRQIAPGQTGKIADYNYKLPDDFTGCVIAWNSNKLDSKVG GNYNYLYRLFRKSNLKPFEKDISTEIYQAGSKPCNGVEGFNCYKPLQSYGFQPTNGVGYQPYRV VVLSFELKHAPATVCGPKKSTNLG<u>HHHHHH</u>

#### R11K

RVQPTESIVRFPNITNLCPFGEVFNATKFASVYAWNRKRISNCVADFSKLYNSASFSTFKCYGV SPTKLNDLCWTNIYADSFVIRGDEVRQIAPGQTGKIADYNYKLPDDFTGCVIAWNSNKLDSKVG GNYNYKYRLFRKSNLKPFEKDISTEIYQAGSKPCNGKEGFNCYKPLQSYGFKPTNGVGYQPYRV VVLSFELKHAPKTVCGPKKSTNLG<u>HHHHHH</u>

#### R12K

RVQPTESIVRFPNITKLCPFGEVFNATKFASVYAWNRKRISNCVADFSKLYNSASFSTFKCYGV SPTKLNDLCWTNIYADSFVIRGDEVRQIAPGQTGKIADYNYKLPDDFTGCVIAWNSNKLDSKVG GNYNYKYRLFRKSNLKPFEKDISTEIYQAGSKPCNGKEGFNCYKPLQSYGFKPTNGVGYQPYRV VVLSFELKHAPKTVCGPKKSTNLG<u>HHHHHH</u>

#### ACE2-Fc

STIEEQAKTFLDKFNHEAEDLFYQSSLASWNYNTNITEENVQNMNNAGDKWSAFLKEQSTLAQM YPLQEIQNLTVKLQLQALQQNGSSVLSEDKSKRLNTILNTMSTIYSTGKVCNPDNPQECLLLEP GLNEIMANSLDYNERLWAWESWRSEVGKQLRPLYEEYVVLKNEMARANHYEDYGDYWRGDYEVN GVDGYDYSRGQLIEDVEHTFEEIKPLYEHLHAYVRAKLMNAYPSYISPIGCLPAHLLGDMWGRF WTNLYSLTVPFGQKPNIDVTDAMVDQAWDAQRIFKEAEKFFVSVGLPNMTQGFWENSMLTDPGN VQKAVCHPTAWDLGKGDFRILMCTKVTMDDFLTAHHEMGHIQYDMAYAAQPFLLRNGANEGFHE AVGEIMSLSAATPKHLKSIGLLSPDFQEDNETEINFLLKQALTIVGTLPFTYMLEKWRWMVFKG EIPKDQWMKKWWEMKREIVGVVEPVPHDETYCDPASLFHVSNDYSFIRYYTRTLYQFQFQEALC QAAKHEGPLHKCDISNSTEAGQKLFNMLRLGKSEPWTLALENVVGAKNMNVRPLLNYFEPLFTW LKDQNKNSFVGWSTDWSPYAD***ENLYFQG***GSGG*DKTHTCPPCPAPELLGGPSVFLFPPKPKDTLM ISRTPEVTCVVVDVSHEDPEVKFNWYVDGVEVHNAKTKPREEQYNSTYRVVSVLTVLHQDWLNG KEYKCKVSNKALPAPIEKTISKAKGQPREPQVYTLPPSRDELTKNQVSLTCLVKGFYPSDIAVE WESNGQPENNYKTTPPVLDSDGSFFLYSKLTVDKSRWQQGNVFSCSVMHEALHNHYTQKSLSLS PGK*

#### R-KO4

RVQPTESIVRFPNITNLCPFGEVFNATRFASVYAWNRKRISNCVADFSVLANSESFSHFNCYGV EPYKLNDLCFTNVYADSFVIRGDEVAQIAPGQTGKIADYNYKLPDDFTGCVIAWNSNNLDSKVG GNYNYLYRLFRKSNLKPFERDISTEIYQAGSTPCNGVEGFNCYFPLQSYGFQPTNDVDYQPYRV VVLSFELLHAPATVCGPKKSTNLG<u>HHHHHH</u>G**GLNDIFEAQKIEWHE**

